# Integrated Single-Cell Analysis of Multicellular Immune Dynamics during Hyper-Acute HIV-1 Infection

**DOI:** 10.1101/654814

**Authors:** Samuel W. Kazer, Toby P. Aicher, Daniel M. Muema, Shaina L. Carroll, Jose Ordovas-Montanes, Carly G. K. Ziegler, Sarah K. Nyquist, Emily B. Wong, Nasreen Ismail, Mary Dong, Amber Moodley, Krista L. Dong, Zaza M. Ndhlovu, Thumbi Ndung’u, Bruce D. Walker, Alex K. Shalek

## Abstract

Cellular immunity is critical for controlling intracellular pathogens, but the dynamics and cooperativity of the evolving host response to infection are not well defined. Here, we apply single-cell RNA-sequencing to longitudinally profile pre- and immediately post-HIV infection peripheral immune responses of multiple cell types in four untreated individuals. Onset of viremia induces a strong transcriptional interferon response integrated across most cell types, with subsequent pro-inflammatory T cell differentiation, monocyte MHC-II upregulation, and cytolytic killing. With longitudinal sampling, we nominate key intra- and extracellular drivers that induce these programs, and assign their multi-cellular targets, temporal ordering, and duration in acute infection. Two individuals studied developed spontaneous viral control, associated with initial elevated frequencies of proliferating cytotoxic cells, inclusive of a previously unappreciated proliferating natural killer (NK) cell subset. Our study presents a unified framework for characterizing immune evolution during a persistent human viral infection at single-cell resolution, and highlights programs that may drive response coordination and influence clinical trajectory.

## Introduction

Understanding the dynamics of host-pathogen interactions during acute viral infection in humans has been hindered by limited sample availability and technical complications associated with comprehensively profiling heterogeneous cellular ensembles. To date, microarray and bulk transcriptomic studies of yellow fever vaccination^1^ and influenza infection^2^ have highlighted complex cellular responses that vary as a function of time, largely characterizing a common systemic interferon stimulated gene (ISG) program. In each instance, additional insights might be gleaned through more sensitive, discretized systems-approaches that can elucidate the contributions of individual cellular components and nominate features that drive productive responses essential to improve vaccines.

Recently, high-throughput single-cell RNA-sequencing (scRNA-Seq) has emerged as a powerful tool to characterize, transcriptome-wide, complex human systems in health and disease at single-cell resolution^3–9^. When applied to a collection of samples across a disease setting, this approach provides a platform for investigating cell types, states, interactions, and drivers associated with that disease; this information can be used to develop testable hypotheses on therapeutic modulations that may ameliorate disease state^7,8^. Meanwhile, within an individual, longitudinal sampling provides an opportunity to decipher, at unprecedented resolution and absent potentially confounding inter-individual variability^7^, shifts in these same variables, and to associate observed changes with internal or external perturbations^10–12^. Such sampling of a host’s exposure to a pathogen could provide foundational insights into essential cellular response features and their coordination, empowering the rational design of improved prophylactic interventions.

Illustratively, a better understanding of the interplay between innate and adaptive immune responses at the very earliest stages of a viral infection, and its impact on long-term disease, could reveal principles to accelerate prevention efforts. Human Immunodeficiency Virus (HIV) has been the subject of thorough study, and thus is a well-considered model system for examining host responses to a pathogen. Moreover, although the development of antiretroviral therapy (ART)^13^, as well as implementation of pre-exposure prophylaxis (PrEP)^14^ and combination prevention efforts, has improved the lives of persons living with HIV, increased life expectancies, and reduced the number of new infections, there were still 2 million new cases of HIV-1 infection in 2017^15^. This highlights a pressing need for effective HIV vaccines informed by an understanding of natural host-pathogen dynamics.

Here, we apply scRNA-Seq to perform an integrated longitudinal analysis of implicated cell programs and drivers during the critical earliest stages of HIV infection. By examining individuals in the Females Rising through Education, Support and Health (FRESH) study^16,17^ – a unique prospective cohort of uninfected young women at high risk of contracting HIV who are monitored for acute viremia by twice weekly plasma sampling – and focusing on those who were enrolled at a time when standard of care did not include treatment during acute disease, we comprehensively examine untreated cellular immune dynamics during the evolution of hyper-acute infection into chronic viremia. Among over 65,000 cells obtained from repeated sampling of peripheral blood, we identify cell types, states, gene modules, and molecular drivers associated with coordinated immune responses to a viral pathogen. Further, these data suggest candidate cellular features that may influence the magnitude of chronic viremia, known to predict long-term infection outcome. Overall, our longitudinal, granular approach captures multiple dynamic and coordinated immune responses – both shared and distinct between cell types and individuals – and provides a framework for their elucidation in health and disease.

## Results

### Longitudinal single-cell transcriptomic profiling captures major and granular immune subsets in hyper-acute infection

In order to globally and longitudinally examine host immune responses during a hyper-acute infection, we performed scRNA-Seq on peripheral blood mononuclear cells (PBMCs) from four individuals enrolled in FRESH who became infected with HIV, assessing multiple timepoints from pre-infection through one year following initial detection of viremia (Fig. 1A, table S1). In our study, hyper-acute infection refers to timepoints at and prior to peak-viral load, whereas acute infection refers to timepoints after peak viral load but before 6 months. Samples were processed in duplicate using Seq-Well^18^ – a portable, low-input massively-parallel scRNA-Seq platform designed for clinical specimens – allowing for robust single-cell transcriptional analysis of PBMC subsets. All individuals studied demonstrated the expected rapid rise in plasma viremia and drop in CD4+ T cell counts that typify hyper-acute and acute HIV infection (Fig. 1B). Among all individuals, we captured 65,842 cells after eliminating low quality cells and multiplets (see Methods), with an average of 2,195 cells per individual per timepoint. Alignment to a combined human and HIV genome at peak infection timepoints yielded few reads mapping to HIV; therefore, alignment for all samples was conducted using a human-only reference.

**Fig. 1:**
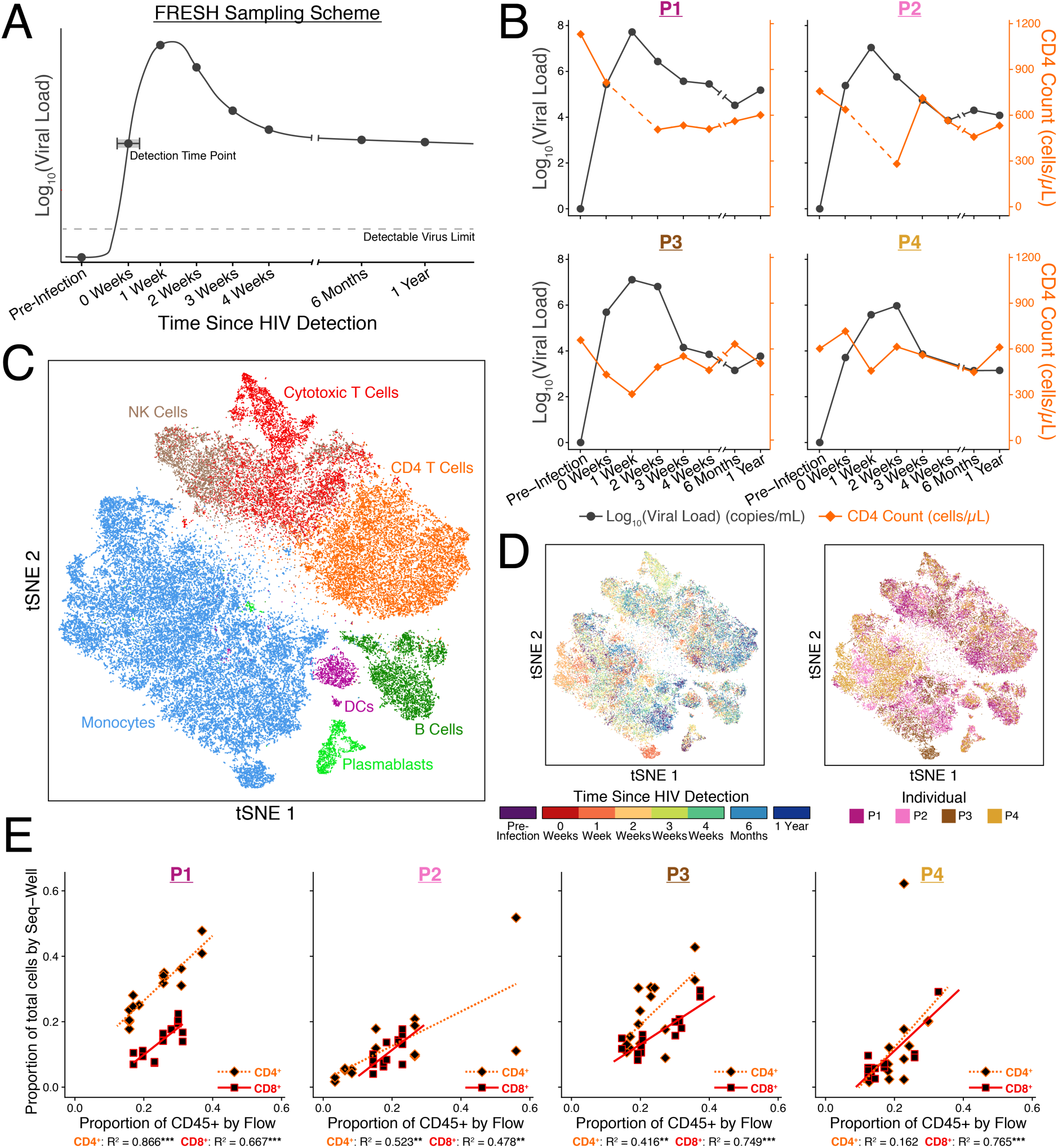
Longitudinal profiling of peripheral immune cells in hyper-acute and acute HIV-infection by single-cell RNA-sequencing. (**A**) Representation of the typical trajectory of HIV viral load in the plasma during hyper-acute and acute HIV infection, and the timepoints sampled in this study. Since participants are tested twice weekly, there is an uncertainty of up to 3 days in where on the viral load curve the first detectable viremia occurs. The exact days sampled are available in table S1. (**B**) Viral load and CD4 T cell count for the four individuals assayed in this study. Dotted lines indicate a missing data point for the metric. (**C**) tSNE analysis of PBMCs from all individuals and timepoints sampled (n=65,842). Cells are annotated based on differential expression analysis on orthogonally discovered clusters. (**D**) tSNE in **C** annotated by timepoint (left) and individual (right). (**E**) Scatter plot depicting the correlation between cell frequencies of CD4+ and CD8+ T cells measured by Seq-Well and FACS. R-squared values reflect variance described by a linear model. * p < 0.05; ** p < 0.01; *** p < 0.001.

To assign cellular identity, we performed variable gene selection, dimensionality reduction, clustering, and embedding *en masse* across data collected from all individuals and timepoints (see Methods). Samples were combined for cell type/phenotype identification to find common transcriptional features of ubiquitous cell subsets, and to improve statistical power on classifying small/rare cell types. Importantly, combined analyses yielded few individual-specific features in the resulting clustering and embedding, suggesting that disease biology, rather than technical batch, is the main driver of variation and subsequent clustering (Fig. 1D, Fig. S1A,B). We annotated identified clusters by comparing differentially expressed (DE) genes that defined each to known lineage markers and previously published scRNA-Seq datasets^19–21^ (Fig. S1C, see Table S2 for list of DE markers). These clusters recapitulated several well-annotated PBMC subsets (Fig. 1C), in addition to revealing phenotypic groupings of monocytes (anti-viral, inflammatory, non-classical) and cytotoxic T cells (CD8+ CTL, proliferating; see Fig. S1D). Thus, we readily and reproducibly mapped the cellular players and phenotypes present along the course of disease progression.

### Cell frequency over time is readily obtained from transcript-assigned cellular identity

We next examined cellular dynamics over the course of infection, beginning with a pre-infection time point. Onset of HIV infection is typically accompanied by an initial depletion of CD4^+^ T cells in the blood and a subsequent small rebound before continued depletion in chronic infection^22^. To ensure that our estimated frequencies would recapitulate conventional measurements of our samples, in parallel, we employed flow cytometry to independently establish the frequencies of T cell subsets (Fig. S2A). Linear regression of the measured CD3^+^CD4^+^ and CD3^+^CD8^+^ flow populations (% of total CD45^+^ cells) with their respective single-cell transcriptome clusters (% of total single cells) across time yielded strong correlations (linear regression, F-test): average CD4^+^ – R^2^ = 0.491, p = 0.0416; average CD8^+^ – R^2^ = 0.665, p = 0.00158 (Fig. 1E). Subsequently, we calculated frequencies for the other cell types in our scRNA-Seq data as a function of time (Fig. S2B). In each individual, we measured an expansion in monocytes at HIV detection and in NK cells that peaked at 3- or 4-weeks post-detection, in-line with studies of influenza and murine cytomegalovirus (MCMV) demonstrating expansion and recruitment of monocytes and NK cells to sites of infection, though on shorter time-scales^23–25^. Altogether, our data elucidate dynamic temporal shifts in the abundance of different cellular subsets during hyper-acute and acute HIV infection aligned with flow cytometry; more importantly, with whole transcriptome information, they enable further global characterization of subcellular activity within and between these subsets.

### Discovering structured variation in cell phenotypes over time in response to infection

To understand how the identified cell types – monocytes, dendritic cells (DCs), plasmablasts, B cells, natural killer (NK) cells, CD4^+^ T cells, CD8^+^ T cells, and proliferating T cells (a sub-cluster of CTLs, see Fig S1D) – varied in phenotype over the course of infection, we assessed coordinated changes in gene expression within each cell type that significantly varied in time. Since the immune responses and time courses of infection were heterogeneous among participants due to our sampling scheme and natural human variability, we performed analyses on an individual-by-individual and cell-type-by-cell-type basis. In this way, our results are sensitive to both intra- and inter-individual changes in gene expression.

To identify tightly co-regulated modules (M) of genes for each type for each individual, we applied weighted gene correlation network analysis (WGCNA)^26,27^ on all cells classified as a particular cell type across all timepoints (Fig. 2A; see Methods for details). Strongly correlated gene modules (permutation test for within-module similarity, FDR corrected q < 0.05) were then tested for significant variation over time by scoring cells at each timepoint against the genes within a module, followed by tests for shifts in score distribution between pairs of timepoints (Wilcoxon rank sum test, FDR corrected q < 0.05). This generated 0-8 temporal modules per cell type (for a list of all significant modules see Table S3 for gene membership and Table S4 for median module scores over time).

**Fig. 2:**
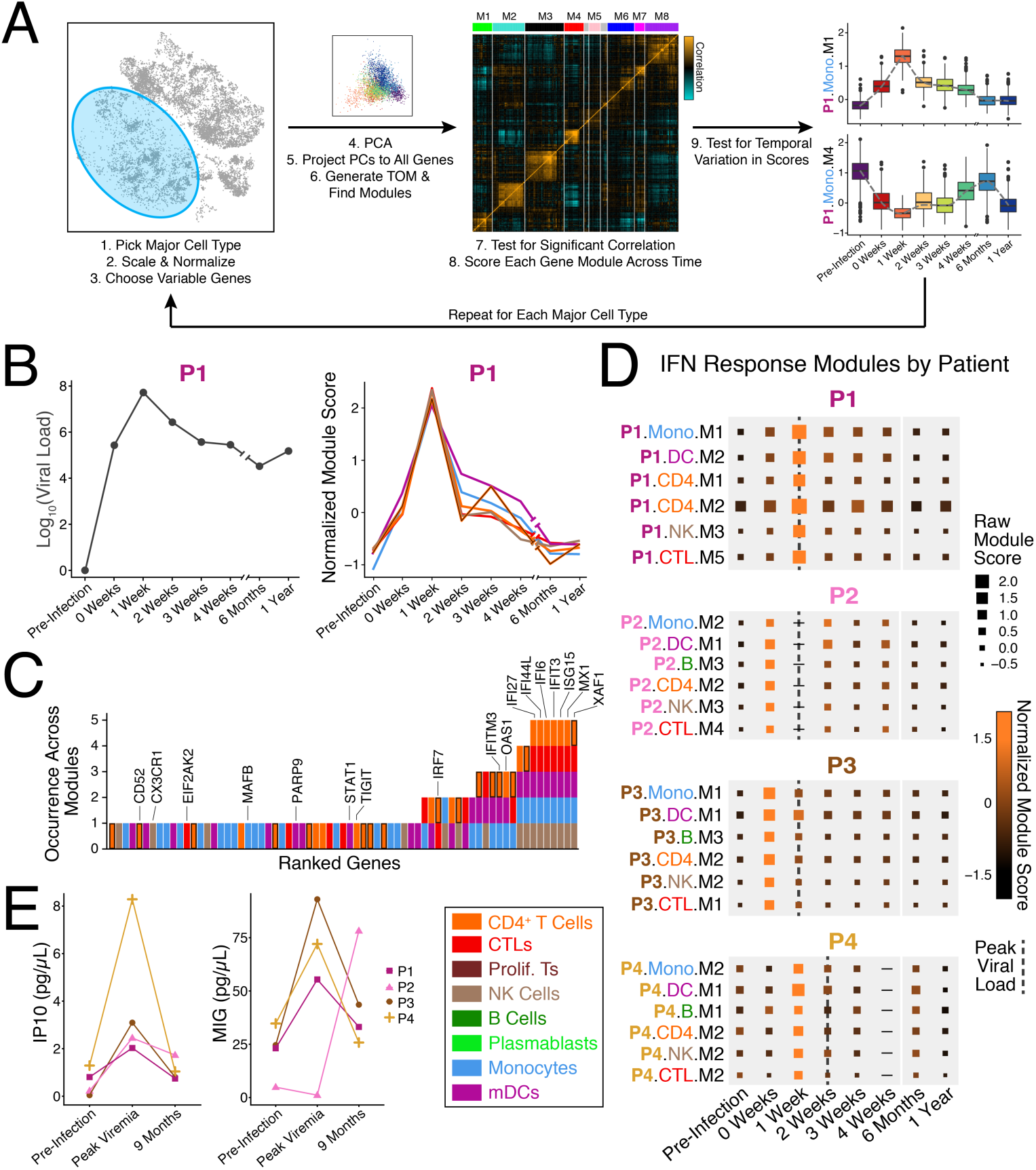
Gene module discovery reveals ubiquitous response to interferon with cell type specific features. (**A**) Schema depicting temporal gene module discovery (see **Methods**). This procedure is repeated for each major cell type (monocytes, CD4+ T cells, CTLs, proliferating T cells, NK cells, B cells, plasmablasts, and mDCs) on an individual-by-individual basis. (**B**) In P1, six gene modules across multiple cell types exhibit similar temporal profiles with peak module scores at the same timepoint as peak viremia is measured. (**C**) Number of occurrences of genes across the modules in **B**. (**D**) Module scores for interferon response modules in each individual. The timepoint where peak viral load occurs is indicated by a dotted line. (**E**) Luminex measurements of IP10 (left) and MIG (right) in matching plasma samples. Points are averages of duplicate measurements.

Across cell types within an individual, these gene program trajectories demonstrated common transient patterns along the course of infection, indicating the utility of this approach in identifying groups of genes acting in concert. While a similar approach is possible using bulk RNA-seq data, here, we are powered to identify temporally similar modules active in distinct subsets of cells both within and across time. Compared to a directed approach, this discovery-based identification of temporally-variant modules enables unbiased selection of coordinated genes and pathways, and immediately reveals differences in response dynamics among cell types, states, and individuals.

### Temporal module analysis reveals shared and unique responses to interferon across cell subsets near peak viremia

With distinct, temporally-variant modules across all cell types and individuals in hand, we next sought to understand these response modules and their association with plasma viral load, the main clinical parameter linked with disease progression rate and clinical outcome^28,29^. Beginning with one individual (P1), we identified a set of 6 significant gene modules spanning multiple cell types that all shared their highest relative module score at the peak viremia timepoint (Fig. 2B). Upon inspection of the genes within each, we uncovered a core set of genes shared among the modules from all cell types: *IFI27, IFI44L, IFI6, IFIT3, ISG15,* and *XAF1*. These genes, in addition to many others belonging to one or multiple of these peak viral load modules, are all induced by type I interferon (IFN-I) stimulation in cell lines and *ex-vivo* primary cells^30–32^ (Fig. 2C, Fig. S3A). Since these modules were generated *de-novo*, our results also report cell type specific genes and functions that correlate with the core measured IFN response signature: anti-viral activity (*CXCL10, DEFB1, IFI27L1*) in monocytes^33,34^, DC activation (*PARP9, STAT1*) likely through sensing of HIV by pattern recognition receptors and interferon by interferon receptors ^35–37^, differentiation of naïve CD4+ T cells (*CD52, TIGIT, TRAC*) potentially into HIV-specific T helper cells^38–41^, and NK cell trafficking (*CX3CR1, ICAM2*) shown to occur in other viral infections^42–44^.

As transcriptional work in humans has been limited to late-acute stage and treated infection^45^, we sought to contextualize our data against the massive IFN response measured in acute SIV infection^46–49^, specifically in rhesus macaques (RM, see Fig. S3B)^47^. In SIV models, natural hosts of the infecting virus (e.g., sooty mangabeys) resolve IFN immune activation more quickly than susceptible hosts, positing that time to resolution may reflect future control in chronic infection (>180 days). By comparison, we find that many IFN stimulated genes induced in RM for 2+ weeks arise and resolve within one week (i.e., upregulate at one timepoint). Here, we are powered to assign the cells expressing these various response genes. For example, upregulation of RIG-I (*DDX58*) is limited to myeloid cells – though RIG-I signaling has been shown to be subverted by HIV^50^ – whereas only CD4^+^ T cells exhibit higher levels of *STAT2*, suggesting a polarization towards a T_H_1 phenotype^51^.

Subsequently, we examined the expression of *IRF7*, one of the interferon regulatory factors that is responsible for anti-viral mediated IFN-I production in SIV/HIV^52,53^ and other viral infections, to determine which cells might be generating this pervasive wave of IFN. In individual P1, almost all cell types demonstrated higher expression of *IRF7* compared to pre-infection and 1-year timepoints (Fig. S3C), highlighting the pervasiveness of IFN-I in response to high levels of viremia and potentially indicative of the positive feedback loop it induces^54–56^. Since plasmacytoid DCs (pDCs) are known to produce IFN-*α* and IFN-*β* in response to HIV^57^, we also assayed single pDCs at peak viremia and 1-year post-infection using a plate-based scRNA-Seq method compatible with enrichment by FACS (Smart-Seq2^58^) (Fig. S4A). At both times, type I IFNs were undetectable (see Supplementary Text). Comparing pDCs between them, we observe modestly increased expression of *IRF7* at peak viremia (Wilcoxon rank sum test, FDR corrected q < 1, log(Fold Change) = 0.42). However, these cells also upregulated several ISGs that were present in the modules of other cell types (Fig. S4B).

We next sought to identify whether similar gene expression programs typified responses in the other three individuals assayed. We readily discovered a similar set of modules around the time of peak viremia in each individual (Fig. 2D and Fig. S3D), as well as shared responses among pDCs (Fig. S4C). Comparing modules across our cohort, we noted common response genes (present in 3 or more cell-types) either shared (*ISG15, IFIT3, XAF1*) or specific (*APOBEC3A, IFI27, STAT1*) to subsets of individuals, suggesting potential core programming and the possibility for the same immune drivers to induce distinct gene responses (Fig. S4D). Finally, to confirm the presence of downstream cytokines from IFN stimulation, we measured MIG (*CXCL9*) and IP10 (*CXCL10*) levels in plasma at pre-infection, peak viremia, and 9-months post infection (Fig. 2E; Methods). All four individuals demonstrated higher levels of IP10 at peak viremia, and three demonstrated elevated levels of MIG. Together, these data highlight the ability of our approach to ascertain a short, pervasive wave of IFN responses in most peripheral immune cells that coincides with, or precedes, peak viremia in hyper-acute HIV infection. Moreover, we uncover nuanced differences among individuals and cellular subsets in this response, as might be expected for an infection associated with diverse clinical courses (e.g., differences in plasma viremia; Fig. 1B).

### Individuals demonstrate diverse, yet coordinated, immune responses during the first month of infection

To investigate other groups of temporally similar modules, we next applied fuzzy c-means clustering^59,60^ to the median module scores at each timepoint across all cell types on an individual-by-individual basis to generate clusters of modules, hereafter referred to as meta-modules (MMs). We subsequently grouped these MMs by temporal shape (Fig. S5 and see Methods for choice of c). MMs represent gene programming in distinct cell types that demonstrate coordinated temporal patterns – here, various cell-types responding simultaneously to infection – enabling us to link discrete transcriptional responses to their propagators. In addition to the aforementioned MM that contained the majority of the IFN response modules (labeled MM3), the only other MM that spanned the majority of cell types was one enriched for ribosomal protein coding genes (labeled MM5, see table S3) – known to indicate cellular quiescence^61^. MM5 demonstrated temporal profiles defined by minimum module scores (i.e., significantly downregulated) around peak viremia, anti-concordant with the immune activation (i.e., significant upregulation) seen in MM3.

Another MM that shared similar temporal immune responses across individuals was MM1, comprised of responses sustained throughout one-month post-detection. In at least 2 of the 4 individuals studied, we identified sustained response modules with shared genes in CD4^+^ T cells, monocytes, NK cells, CTLs, and proliferating T cells (Fig. 3A-E, see table S5 for overlapping genes). While DCs and B cells also expressed multiple modules within this MM, some modules had low MM membership scores and were excluded (membership < 0.25, labeled with **†** in Fig. S5) or did not share any genes across individuals (Fig. S6A and Supplementary Text).

**Fig. 3:**
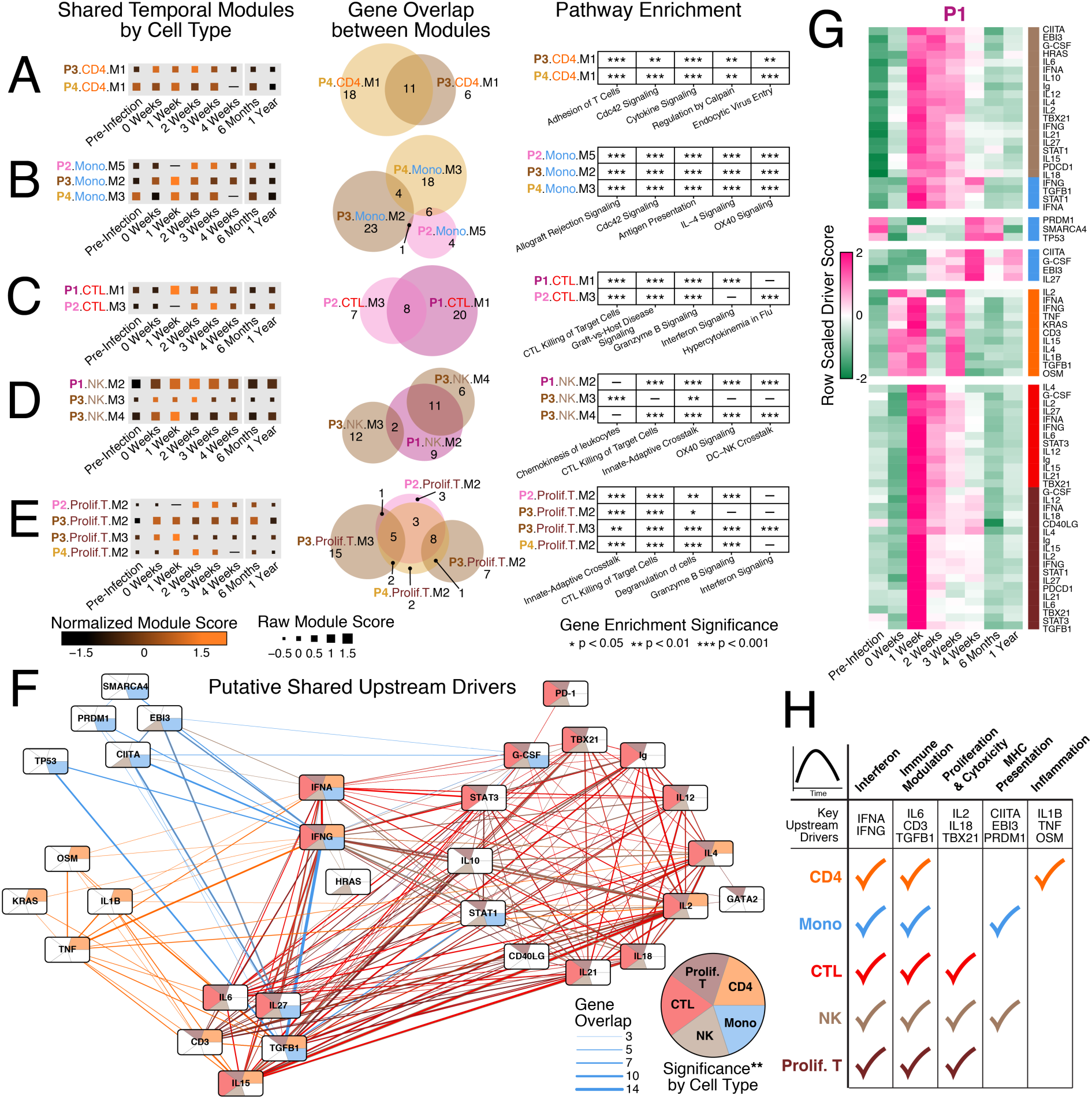
Modules with sustained expression conserved among individuals suggest shared and cell type specific drivers of immune response. Module Scores (left), gene overlaps between modules (middle), and enriched pathways for each module (right) in (**A**) CD4+ T cells, (**B**) monocytes, (**C**) CTLs, (**D**) NK cells, and (**E**) proliferating T cells. (**F**) Network of predicted upstream drivers of modules in **A-E**. Nodes are colored by significance in each cell-type. Edge width and color reflect the number of shared genes (width) in the gene sets of the upstream drivers for a given cell-type (color; see **Methods**). (**G**) Median gene set scores for significantly temporally variant (p < 0.05) upstream drivers in P1. Scores are grouped by k-means clustering; k=5. (**H**) Summary table of immune responses to related and distinct stimuli with similar temporal dynamics.

As each module within MM1 is distinct, we performed gene set enrichment analyses (see Methods) to discern if, in addition to sharing genes, modules from the same cell type shared functional annotations across individuals (Fig. 3A-E). In every cell type, modules across individuals were significantly enriched for many of the same underlying pathways (see table S6 for full list), despite slightly variable temporal dynamics and unique gene membership. CD4^+^ T cells expressed genes associated with non-classical viral entry by endocytosis^62^ as well as adhesion, potentially suggesting migration and viral dissemination throughout the body. Monocytes expressed genes associated with antigen presentation and IL-4 signaling (mainly HLA-DR subunits), which may reflect generalized interferon responses, or the potential to promote active T helper and CTL responses. NK cells, CTLs, and proliferating T cells all upregulated genes associated with killing of target cells by perforin and granzyme release, highlighting the joint role of innate and adaptive cells in combating viremia (see Table S5 and Fig. S6B for all shared responses across cell types)^63,64^. Thus, our results indicate common functional enrichments supported by gene sets that vary across cell types and individuals in response to infection.

### Distinct cell types respond to common and unique upstream drivers induced in infection

To identify common and cell-type specific inducers of these measured transient responses extending past peak viremia, we generated a list of predicted upstream drivers of each module (see Table S6). Selecting highly significant hits in at least two modules, we drew a network of putative upstream drivers (nodes) colored by significance in each cell type with edges connecting nodes with shared enriched genes (Fig. 3F, Fig. S6C, and see Methods). Strikingly, IFN-*α* and IFN-*γ* are predicted drivers of these sustained responses for all five cell types even though these modules do not contain the typical ISGs; in chronic HIV infection, prolonged IFN-I stimulation has been shown to maintain viral suppression but also blunt other immune functions in a humanized mouse model^65,66^. Matching Luminex data confirmed elevated levels of IP-10 and MIG at one-month post HIV detection (Fig. S6D). IL-15 and IL-2, known to induce T and NK cell proliferation but to lead to defects in chronic infection^67–69^, were enriched as drivers for all lymphocytes explored. However, they also shared enriched genes with several other interleukins, including IL-4, IL-12 (also elevated in plasma, see Fig. S6D), and IL-21. Interestingly, only CD4^+^ T cell modules were enriched for TNF, IL-1B, and OSM, suggesting the directed induction of pro-inflammatory T helper cells^70^. Meanwhile, monocytes and NK cells were enriched for CIITA and EBI3 (a subunit of IL-27), which regulate MHC-II and MHC-I genes, respectively^71,72^.

We also contextualized observed responses to these upstream drivers temporally by re-scoring against enriched genes for each driver. This analysis revealed variable kinetics in the onset, intensity, and length of immune responses across different cell types (Fig. 3G, Fig. S7). We note the following gene-programming upregulation trends in most individuals: CD4+ T cells are active from before peak viremia throughout 3-4 weeks post infection, and CTL and proliferating T cell programs are induced for 2-3 weeks around peak viremia, whereas NK cell and monocyte activity extends throughout the first month of infection.

Based on shared cell-type enrichments, genes, and functions, we summarize the multitude of common immune responses displaying sustained gene expression over the course of first month of HIV infection, and their potential drivers, across individuals (Fig. 3H). While the IFN stimulated gene programs do not extend past hyper-acute infection, our data suggest that persistent IFN activation could manifest in different ways in each cell type, leading long-term to previously shown dysfunction partially mediated by IFN in chronic infection^73^. This analysis also support more complex cytokine interactions – some previously described as synergistic (e.g. IL-2 & IL-18^74^) or antagonistic (e.g. IL-6 & IL-27^75^) – occurring in acute infection, and delineates how they may affect various cell types. Though dozens of cytokines are known to elevate in plasma during acute HIV infection^76^, here we present a putative schematic of which cell types they modulate alongside other extracellular proteins and transcription factors active during this time frame. Furthermore, our analysis establishes a blueprint of multi-cellular responses to several stimuli along the course of hyper-acute and acute infection to be edified by application to other pathogens.

### Two instances of temporally similar modules within a cell type discerned by scRNA-Seq

After discovering temporally variant modules in our dataset, we observed a few sets that demonstrated similar temporal response patterns in a given cell type, but were not combined into a single module by our framework. We thus sought to understand how these modules might be linked by looking across single cells for module co-expression. Here, single-cell expression data are essential to distinguish response circuitry among cells.

The clearest example of multiple modules being co-expressed with the same temporal pattern in the same cell type from our analysis was the NK activated M3 module (*CCL3, CCL4, CD38*) and the cytotoxic M4 module (*PRF1, GZMB, HLA-A*) in P3 (Fig. 3D), both part of MM1. Enrichment analysis demonstrated little overlap between the significant pathways associated with these modules, implying orthogonal biological function. We therefore investigated whether the gene programs for these modules were highly co-expressed in the same single cells and thus co-varied among single cells across time (Fig. S8A). While we did not observe differential simultaneous upregulation of these modules between time points, we found variation in the correlation of cell-scores for the pair as a function of time across single cells, with the strongest correlation one to two weeks after HIV detection (Fig. S8B). Variation in the correlation of M3 and M4 may reflect a synergizing of these gene programs^77^ within NK cells to combat HIV as viremia declines post peak.

In examining MM3 (Fig. S5) – containing the majority of the IFN response modules – we observed that P3 also exhibited a set of temporally similar modules in monocytes (M1 & M3); however, these modules did not variably correlate in expression score as a function of time. Instead, these gene programs were highly co-expressed but only at HIV-detection (Fig. S8C-D). Gene set analysis readily demonstrated that monocyte M1 consisted of IFN response genes, while M3 was enriched for genes associated with inflammation (Fig. S8E). IFN has been shown to stunt the production of pro-inflammatory cytokines in monocytes similar to the phenotype observed in these cells in viremic persons^78,79^, but the co-expression of anti-viral and pro-inflammatory signals in the same single cells has not yet been described to our knowledge. As these module scores are generated independently for each single cell, individual monocytes in this person at the time of HIV detection are simultaneously expressing both anti-viral and inflammatory gene programs. Critically, our longitudinal granular, single-cell approach facilitates the study of variation in gene module correlation and co-upregulation, suggesting key cellular circuitry, and its coordination, during response to infection.

### One individual who naturally controls infection displays a polyfunctional subset of monocytes at HIV detection

Intrigued by the appearance of these polyfunctional monocytes in one individual, we next explored whether the other individuals assayed developed similar cells after infection. Scoring monocytes from each individual on inflammatory and anti-viral gene lists derived from discovered modules (Fig. S9A), we were unable to identify these polyfunctional monocytes in the other three individuals (Fig. 4A-B, Fig. S9B-C). In fact, looking at structured gene variation in monocytes over time in principal component analysis (PCA) space revealed that the major axis of variation (PC1) in P1 and P2 not only reflected sample timepoint, but also separated monocytes based on their expression of anti-viral and inflammatory genes. In P3 and P4, however, these gene programs contributed to different principal components, suggesting their independence in defining monocyte phenotype.

**Fig. 4:**
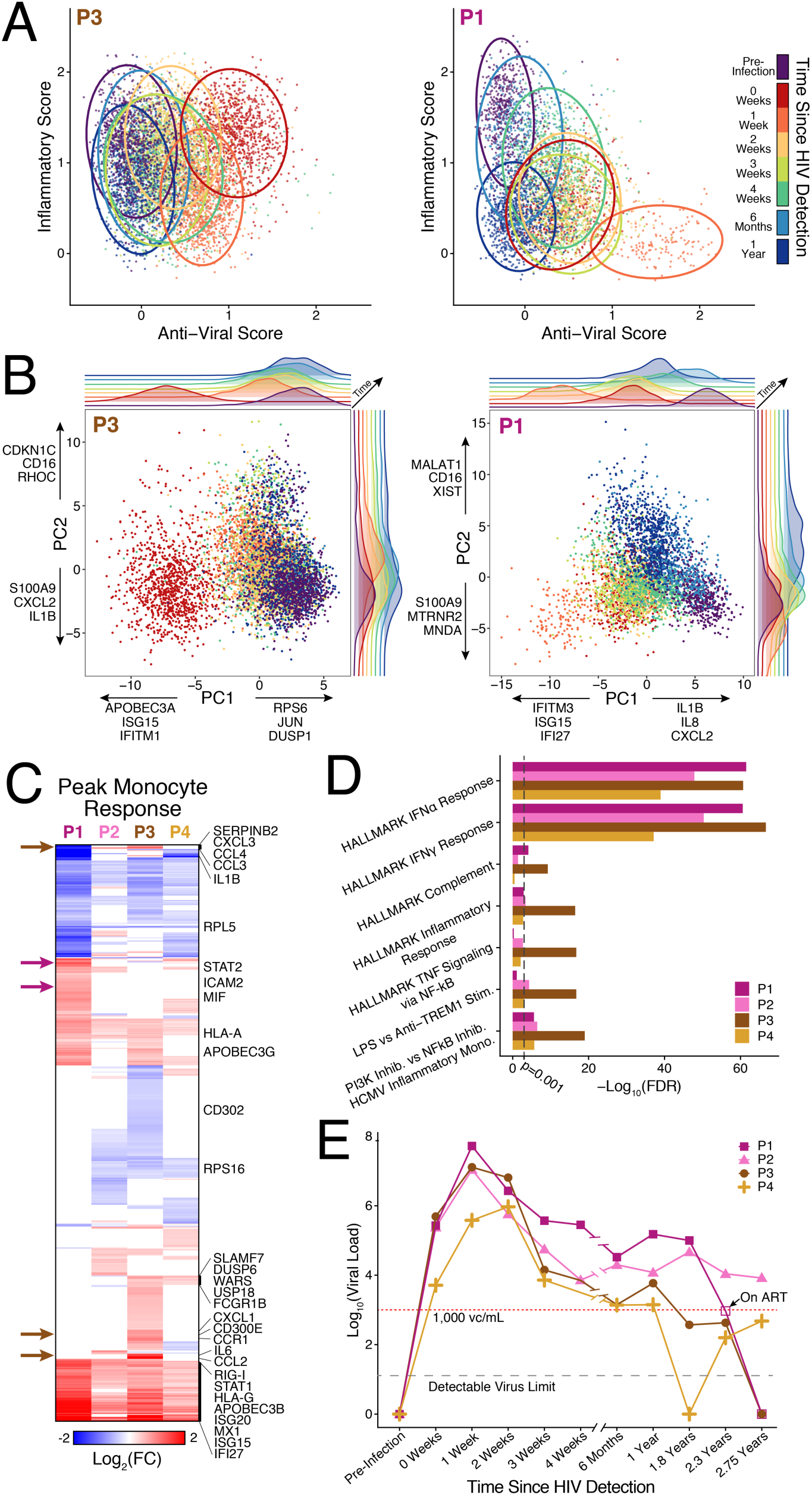
One individual who goes on to control infection presents a poly-functional subset of monocytes at HIV detection. (**A**) Inflammatory and anti-viral scores of monocytes in P3 (left) and P1 (right) derived from gene lists created from merging modules among individuals. Ellipses drawn at 95% confidence interval for cells from each timepoint. (**B**) Principal component analysis (PCA) of all monocytes from P3 (left) and P1 (right). Density of cells in PC1 vs PC2 space annotated by timepoint are depicted, and the top loading genes for PC1 and PC2 are also annotated. (**C**) Heatmap of differentially expressed genes between monocytes at the peak response timepoint (0 weeks/1 week) vs pre-infection. Arrows indicate genes specific to P3 (dark-brown) and P1 (violet). (**D**) Enriched pathways for the differentially expressed genes in **C**, using the MSigDB Hallmark Gene Sets. (**E**) Viral load by RT-PCR of the plasma of the four individuals assayed out to 2.75 years. Controllers of HIV maintain levels of plasma viremia less than 1,000 viral copies (vc)/mL. P1 initiated ART before the 2.3 year timepoint.

In all four individuals, we saw dramatic structuring of the monocytes in PC space by time. Specifically, monocytes sampled at HIV detection (0 weeks) or 1-week post-detection were strongly separated along either PC1 or PC2, indicating a pervasive hyper-acute response to infection. Interestingly, non-classical monocytes (see Fig. S1D and Table S2), which may be more susceptible to infection^80^, displayed disparate temporal dynamics across individuals, even though they drove significant variation in PCA space (Fig. S9D). Comparing DE genes at these peak response timepoints (vs. pre-infection) highlighted not only the specificity of the co-inflammatory/anti-viral monocytes to P3, but also other person specific differences in monocyte phenotype (Fig. 4C). Gene set analysis on upregulated genes in each individual confirmed that monocytes in all individuals produced strong anti-viral factors (e.g., *RIG-I, APOBEC3B, MX1*) with significant enrichment (MHC hypergeometric test, q<0.001) for response to IFN-*α* and IFN-*γ* (Fig. 4D). Moreover, corroborating the scoring on inflammatory genes, only P2 and P3 were significantly enriched for inflammatory responses, and only P3 for TNF signaling via NF-kB (MHC hypergeometric test, q<0.001). In fact, P1 and P2 demonstrated downregulation of genes associated with inflammation compared to pre-infection.

Subsequently, we investigated known clinical parameters in our cohort for features of infection that might be related to the appearance of these polyfunctional cells. As the level of viral load in chronic infection correlates with disease outcome^28^, we compared the viral load setpoints of these individuals at 1.8, 2.3, and 2.75 years after HIV detection. Two of the four individuals (P3 & P4) maintained low levels of viremia (< 1,000 viral copies (vc)/mL) out to 2.75 years in the absence of ART (Fig. 4E). HIV infected persons who naturally maintain low levels of viremia in chronic infection (controllers) have been shown to have enhanced immune responses in chronic infection^7,81,82^. However, whether early events in acute HIV infection reflect or contribute to long-term control is unknown. In the hyper-acute monocyte responses (Fig. 4C), we found a small set of upregulated genes shared only by P3 and P4, including *SLAMF7*, whose activation was recently described to downregulate CCR5 on monocytes and reduced their infection capacity by HIV^83^, suggesting a potential difference in monocyte susceptibility and phenotype in these individuals during hyper-acute infection. Moreover, referring back to the initial cell type clustering of our data (Fig. S1), we noted that the peak response monocytes in P3 (0 weeks) clustered separately from other monocytes, and that P4 made up >75% of the anti-viral monocytes detected at 1-week post-infection. Identifying a potential correlate of future viral control otherwise obscured by bulk transcriptomics and sparse longitudinal sampling, we next searched for other unique immune responses enriched in either or both of the two controllers.

### Future controllers exhibit higher frequencies of proliferating CTLs and a precocious subset of NK cells before traditional HIV-specific CD8+ T cells

As CD8+ T cells are known to play a part in controlling chronic HIV infection^82,84,85^, we turned to the CTLs in our study to look for differences between the individuals who controlled infection long-term and those who did not. Through our module discovery approach, we found that CTLs produced increasing levels of *PRF1* and *GZMB* along the course of hyper-acute infection (Fig. 3C). Further unsupervised and directed approaches did not elucidate meaningful or significant differences in CTL responses across individuals by outcome of viral control (Fig. S10A-B and Table S7).

Recently, we demonstrated that, in most individuals in the FRESH study, a majority of proliferating CTLs in hyper-acute infection are HIV-specific by tetramer staining^86^. Therefore, we turned to the proliferating T cells in our study to look for differences in response based on long-term viral control. *En masse*, the proliferating T cells expressed similar levels of cytotoxic genes as non-proliferating CTLs (Fig. S10C). DE analysis highlighted genes associated with cell-cycle (e.g. *STMN1, HIST1H1B, MKI67*) and memory (e.g. *IL7R, KLRB1*) (see Fig. S10D and table S7) for proliferating and non-proliferating CTLs, respectively. While sparsely detected due to the method of library construction in Seq-Well, we did measure a limited number of TCR variable genes in the proliferating CTLs (Fig S10E). In fact, we note enrichment of TRBV and TRAV genes known to construct prevalent CDR3 sequences that bind common HIV epitopes^87,88^: *TRBV28* (QW9/FL8/KF11/KK10/NV9, *χ*^2^ test p=2.4*10^-26^), *TRAV4* (KK10, *χ*^2^ test p=3.5*10^-6^), and *TRBV20-1* (KK10/KF11/GY9/NV9, *χ*^2^ test p=0.059). Our single-cell data here expand our recently published bulk RNA-Seq data on HIV-specific CTLs in this cohort^89^, but also enable us to elucidate heterogeneity in this proliferating cytotoxic response as a function of time.

Grouping proliferating T cells with the other CTLs, we sought to understand if these two controllers demonstrated differences in the frequency of proliferating T cells among the total CTL pool over time. Strikingly, both controllers (P3 & P4) displayed much higher frequencies of proliferating T cells within the first month of infection (Fig. 5A). While all four individuals developed proliferating T cells at 1-week post HIV detection, P3 and P4 exhibited a higher fraction of these cells 1 week after HIV detection (30-40%).

**Fig. 5:**
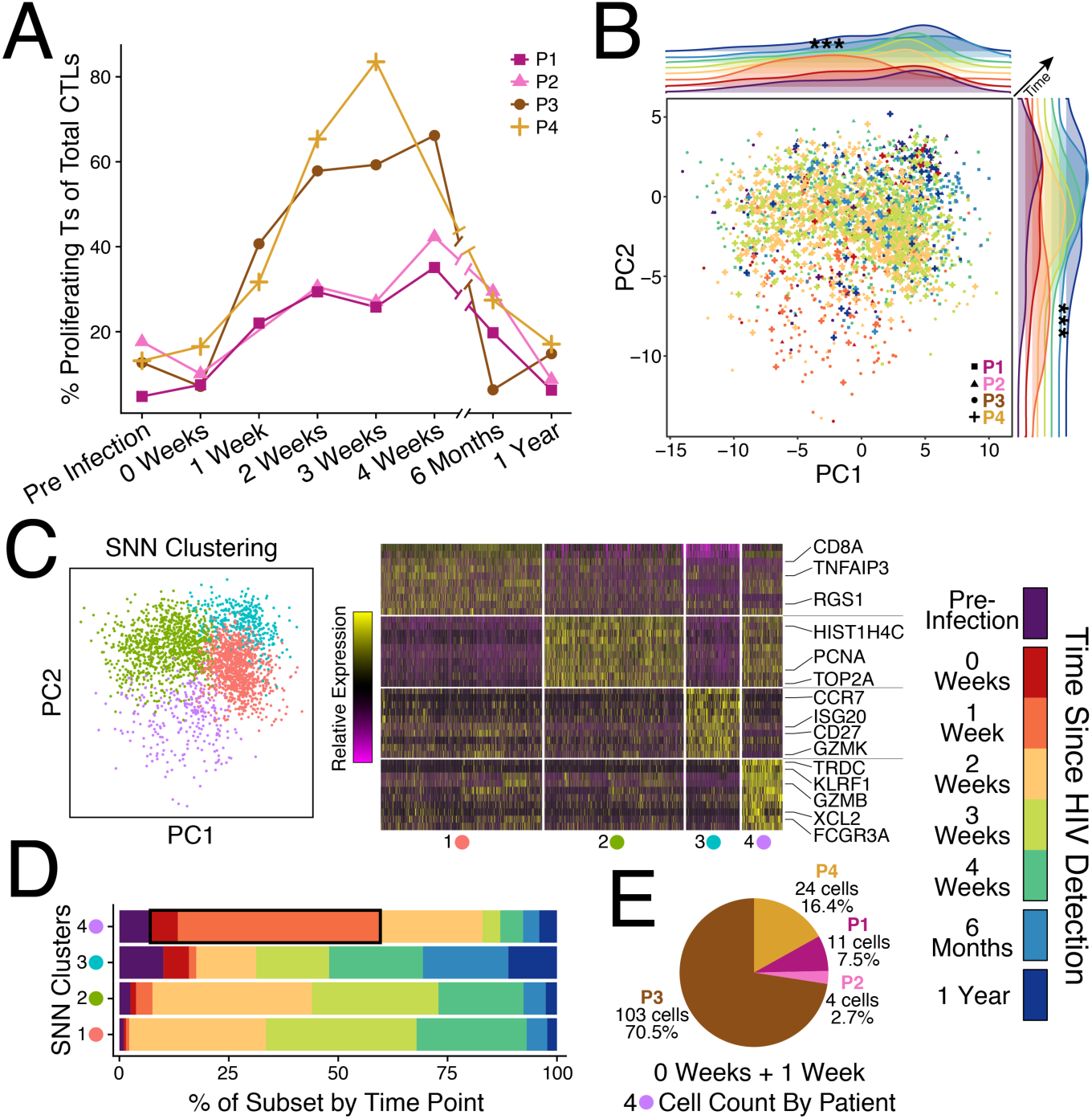
Future controllers exhibit higher frequencies of proliferating CTLs and a precocious subset of NK cells 1 week after detection of HIV viremia. (**A**) Proportion of proliferating T cells of total CTLs as a function of time and individual measured by Seq-Well. (**B**) PCA of proliferating T cells from all four individuals. Cells assayed from the 1-week timepoint strongly separate along PC1 and PC2; Mann Whitney-U Test, *** p < 0.001. (**C**) SNN clustering over the top 6 PCs reveals four sub-clusters (left) with distinct gene programs (right). (**D**) Percentage of cells in each sub-cluster by timepoint. (**E**) Number of cells from each individual within the cells sampled at 0 weeks and 1 week in the NK cell cluster (4-lilac; black box in **D**).

We next utilized unsupervised analyses to explore differences in proliferating T cell responses over time among individuals (Fig. 5B, Fig. S10F). Proliferating T cells captured at 1-week post-infection strongly separated in PCA across both PC1 and PC2 (p < 0.001). Clustering over all proliferating T cells (see Methods), we identified four clusters of cells with distinct gene programs (see Fig. 5C and table S7): traditional CD8+ T cells (1-red), hyper-proliferative CD8+ T cells (2-green), naïve CD4+ T cells (3-cyan), and a subset of cells that is CD8– but *TRDC+* and *FCGR3A*+ (CD16) (4-lilac). A recent scRNA-Seq study on cytotoxic innate-ness looked at cytotoxic *γδ*T and NK cells in healthy humans, noting basal levels of *TRDC* in both cell-types^21^. To determine whether these *TRDC*^*+*^*CD16*^+^ cells were *γδ*T or NK cells, we scored them, as well as non-proliferating CTLs and NK cells, against gene signatures described in that study (Fig. S10G). Based on score similarity to NK cells, and the relative down-regulation of CD3 compared to the other proliferating T cell subsets (Wilcoxon rank sum test; *CD3D:* log(FC) = −0.895, q = 2.7×10^-42^; *CD3G*: log(FC) = −0.923, q = 8.9×10^-37^), we determine cluster 4 (lilac) to be proliferating NK cells. Looking at the distribution of timepoints within each of these clusters, this NK cluster (4-lilac) contained the highest proportion of cells assayed at HIV detection and 1 week thereafter (Fig. 5D,E). Within these earliest proliferating NK cells, the majority were detected from P3 and P4. Together, these data suggest that individuals who go on control HIV infection without ART exhibit a subset of proliferative, cytotoxic NK cells before the majority of HIV-specific CD8+ T cells arise. Thus, investigating the classically induced cytotoxic cells in viral infection on a single-cell level revealed unappreciated heterogeneity in the anti-viral response, implicating innate immune responses in controlling infection.

## Discussion

Here we have applied both unsupervised and directed approaches to a unique longitudinal human infection data set to characterize conserved immune response dynamics, as well as early cellular events associated with the individuals studied here who go on to control infection without treatment. Sampling prior to and immediately upon HIV infection, we assayed longitudinal PBMC samples in four individuals from a prospective cohort, the FRESH Study^16,17^ using Seq-Well^18^. This systems-level approach revealed parameters shared across all cell types examined (e.g., response to IFN), as well as subtle variations among cellular types and individuals missed in previous bulk studies of infection. Further, it defined cell-type specific responses (e.g., inflammatory induction of CD4+ T cells), and their interaction dynamics following infection. Moreover, leveraging the resolution and high-throughput capability of scRNA-Seq methods, we were able to uncover previously unappreciated cellular features in the PBMCs of two individuals who went on to control infection naturally, including subsets of poly-functional monocytes and proliferating NK cells limited to hyper-acute infection, that may correspond to better infection outcome.

To systematically identify immune cells responding with similar temporal dynamics, we adapted WGCNA^26,27^ (Fig. 2A and see Methods) to discover modules of genes that significantly changed in expression within a given cell type over time. Cellular responses to infection can happen on the order of hours to days; therefore, even with the biweekly HIV testing in the FRESH Study, we anticipated these individuals would not align immune responses in absolute time. After applying our module analysis, the strongest and most pervasive module across cell types and all individuals assayed was the interferon induced anti-viral response (Fig. 2D). While known to be a key factor in controlling HIV replication^30,65^ and the major response in NHP SIV infection models^52,90^, the timing of response and extent to which it pervades all peripheral cell subsets in humans has not yet been described. Of note, both controllers (P3 & P4) exhibited interferon response modules the week before peak viremia, consistent with the faster resolution of interferon response in natural SIV hosts compared to non-natural hosts^46–49^. Moreover, multiple modules from P3 & P4 uniquely contained *APOBEC3A*, shown to restrict HIV infection in myeloid cells^91^, and *IFITM1* and *IFITM3* which can inhibit HIV translation in transfected cells *in-vitro*^92^.

Due to our ability to determine enriched modules within individual cells, we were able to unveil a second layer of regulation, which might otherwise be drowned out by the overwhelming IFN signature (Fig. 3F-H). This highlighted putative upstream drivers that are unique to CD4+ T cells, monocytes, NK cells, or shared amongst many cell types. Downstream genes (many shared) were significantly enriched for many known drivers of lymphocyte proliferation, emphasizing the presence of mounting large cytotoxic responses in more than just HIV-specific CD8^+^ T cells during acute infection. Some of these molecules were also upstream of CD4^+^ T cells, potentially increasing their susceptibility to infection (IL-15)^69^ and inducing maturation (IL-2)^67^ and differentiation (IL-4)^93^. Cell-type specific drivers, like IL-1B & TNF upstream of CD4+ T cells, also suggest T helper subset differentiation during this time frame^70^. However, the functional capacity of CD4^+^ T cells to coordinate productive CD8^+^ T cells during hyper-acute HIV infection has yet to be tested. Though we did not ascribe the relationships between all cell types and their immune modulators, this integrated multi-cellular analysis lays the foundation for future characterization of the complex, dynamic immune responses to an infection. A potential method to pinpoint the effects of the various cytokines produced in acute infection might utilize *in-vitro* assays that couple PBMCs from healthy individuals with and without autologously HIV infected CD4+ T cells.

Empowered by our single-cell resolution and cognizant of the role HIV-specific T cells play in long-term control^82,84,94^, we were intrigued to find not only higher frequencies of proliferating CTLs in P3/P4, but also the presence of a subset of a previously unappreciated proliferating NK cells preceding the well-described HIV-specific responses (Fig. 5C-E), given the multi-faceted role of NK cells in viral control^64^. Assaying cells from controllers *in-vitro* showed that NK cells were equivalent to CD8^+^ T cells in inhibiting viral replication^95^; however recent work has demonstrated CD11b^+^CD57^-^CD161^+^Siglec-7^+^ NK cells to be more abundant in elite controllers compared to those who progress^96^. The proliferating NK cells measured here also express high levels of CD161 (*KLRB1*), associated with the production of IFN-*γ* in response to IL-12 and IL-18^97^. Antigen specific expansion of cytotoxic NK cells has been shown to occur in hCMV^98,99^, hantavirus^100^, and SIV^101^ as a “memory-like” response; however, we do not measure changes in NKG2C (*KLRC1*) here. Lacking the opportunity to assay previous viral exposure in these individuals, we cannot comment on whether these cells might be proliferating in response to a previously encountered antigen from HIV or a similar retrovirus. We hypothesize that a similar phenotype of proliferating NK cells may arise in response to re-encountering antigen after early ART. To test this, one could examine the killing capacity of NK and CD8^+^ T cells *in-vitro* from individuals treated at various stages of acute and chronic infection, given sample availability.

Collectively, our single-cell transcriptional study of hyper-acute and acute HIV infection in FRESH provides several key insights into the dynamics of host-immune responses to infection on a systems-level. It also affords a key reference data set for studying the earliest moments of viral infection after detection. While limited sample availability and the inability to recreate a prospective study like this (since immediate ART is now standard of care) preclude strong associations with clinical parameters across individuals, we are able to nominate potential early responses that may inform long-term viral control and thus guide HIV vaccine efforts. Although preliminary, many of these observations can be validated in NHP models via proper selection of natural and unnatural hosts/virus strains. Future work in FRESH will seek test the effects of early administered ART on these longitudinal HIV response dynamics, while work in other viral and bacterial infections in additional human cohorts will enable assessment of the broad utility of the methods and features described here.

## ACKNOWLEDGEMENTS

We thank the individuals who have participated in the FRESH Study, as well as the FRESH and HIV Pathogenesis Programme (HPP) staff; M. Waring, N. Bonheur, E. Koscher for flow cytometry and sorting services through the Ragon Institute Imaging Core-Flow Cytometry Facility; A. Piechocka-Trocha for sample handling and reagent preparation; M. Cole and N. Yosef for advice on computational methodology; D. Kwon, S. Bloom, M. Hayward for participant intake and STI data; M. Carrington and A. Bashirova for HLA genotyping; S. Rasehlo and N. A. Akilimali for support with Luminex measurements; A. Leslie, H. Kløverpris, D. Lingwood, and S. Pillai for insightful discussions. This work was supported, in part, by the Searle Scholars Program, the Beckman Young Investigator Program, the Pew-Stewart Scholars Program for Cancer Research, a Sloan Fellowship in Chemistry, the NIH (2U19AI089992, 2R01HL095791, 1U54CA217377, 2P01AI039671, 5U24AI118672, 2RM1HG006193, 1R33CA202820, 1R01AI138546, 1R37AI067073, 1R01HL126554, 1R01DA046277, 1U2CCA23319501, K08 AI118538), the Bill and Melinda Gates Foundation (OPP1139972, OPP1202327, OPP1137006, OPP1202327, OPP1066973, OPP1146433), an NSF Graduate Student Fellowship Award, the Hugh Hampton Young Memorial Fund Fellowship, Gilead Sciences, Inc (Grant ID #00406), the International AIDS Vaccine Initiative (IAVI) (UKZNRSA1001), the South African National Research Foundation (grant #64809), the NIAID (R37AI067073), the Witten Family Foundation, Dan and Marjorie Sullivan Foundation, the Mark and Lisa Schwartz Foundation, Ursula Brunner, the AIDS Healthcare Foundation, and the Harvard University Center for AIDS Research (CFAR, P30 AI060354, which is supported by the following institutes and centers co-funded by and participating with the US National Institutes of Health: NIAID, NCI, NICHD, NHLBI, NIDA, NIMH, NIA, FIC and OAR.), the Sub-Saharan African Network for TB/HIV Research Excellence (SANTHE), a DELTAS Africa Initiative [grant # DEL-15-006]. The DELTAS Africa Initiative is an independent funding scheme of the African Academy of Sciences (AAS)’s Alliance for Accelerating Excellence in Science in Africa (AESA) and supported by the New Partnership for Africa’s Development Planning and Coordinating Agency (NEPAD Agency) with funding from the Wellcome Trust (grant # 107752/Z/15/Z) and the United Kingdom (UK) government. The views expressed in this publication are those of the author(s) and not necessarily those of AAS, NEPAD Agency, Wellcome Trust or the UK government. Raw data will be available through Database of Genotypes and Phenotypes (dbGaP), accession number pending.

## SUPPLEMENTARY INFORMATION

Methods, supplementary discussion, and supplementary figures can be found in the Supplementary Information file. Supplementary tables are available upon request from the corresponding author.

