## Supplementary Information for "Integrated Single-Cell Analysis of Multicellular Immune Dynamics during Hyper-Acute HIV-1 Infection"

### MATERIALS AND METHODS

*Study Subjects*

All individuals in this study were participants in the FRESH Cohort[16,17]. This prospective study enrolls HIV negative women, ages 18-24, and tests for HIV-1 RNA in the plasma twice a week for one year. Each time the women come to the study center, they participate in peer-support groups and receive a stipend. In addition to semi-weekly virus testing by RT-PCR, whole blood is collected 4 times (including during enrollment) throughout the year from participants. If a plasma test comes back positive, the participant is asked to come back to the clinic that day to collect a blood sample. Samples are then collected weekly through the first 6 weeks of infection, and regularly afterward as long as the individual continues to return to the study center. In the arm of the study described herein, subjects were initiated on anti-retroviral therapy (ART) when their CD4 count fell below 350 cells/μL, per standard treatment guidelines at the time of enrollment. A second arm of the study was initiated in 2014, and is currently still in place; in it, individuals who test positive for viral RNA are initiated on ART when they are called back into the study center for their first post-infection sample collection. To the best of our knowledge, all individuals in this study had not yet started ART for the time points processed here. FRESH was performed in accordance with protocols approved by the institutional review board at Partners (Massachusetts General Hospital, Boston, USA), MIT (Cambridge, USA) and the biomedical research ethics committee of the University of KwaZulu-Natal (Durban, South Africa).

*Cell preparation, flow cytometry, and cell sorting*

Frozen peripheral blood mononuclear cells (PBMCs) were thawed and washed twice with warm RPMI supplemented with 10% fetal bovine serum. Next, the cells were resuspended in FACS buffer (PBS supplemented with 1% FBS) and stained with antibodies on ice for 30 minutes in FACS buffer. Antibodies used include Alexa Fluor 700 - CD45 (Biolegend, clone 2D1), BUV737 – CD3 (BD Biosciences, clone UCHT1), BV711 – CD4 (Biolegend, clone OKT4), BUV395 – CD8 (BD Biosciences, clone RPA-T8), BV605 – CD14 (Biolegend, clone M5E2), BV510 – HLA-DR (BD Biosciences, clone G46-6), and BV650 – CD123 (Biolegend, clone 6H6); subsets of these markers were used to identify immune cells ($CD45^+$), $CD4^+$ ($CD45^+CD14^-CD3^+CD4^+$) and $CD8^+$ ($CD45^+CD14^-CD3^+CD8^+$) T cells, and pDCs ($CD45^+CD14^-CD3^-CD11c^-HLA\text{-}DR^+CD123^{++}$). Afterward, the cells were washed and stained with the viability stain Calcein Blue, AM (Invitrogen, C34853) for 15 minutes on ice. Finally, the stained cells were washed twice with FACS buffer and sorted on a BD SORP FACSAria II cell sorter using BD FACSDiva software and a 100-micron nozzle. Up to 250,000 viable immune cells ($CD45^{+}$Calcein $Blue^{+}$) were sorted into 1 ml of RPMI + 10% FBS for Seq-Well[18]. For Smart-Seq2[58], individual cells were directly sorted into 10 µl of RLT (Qiagen) + 1% BME in 96 well plates.

*Single-cell RNA-Seq (scRNA-Seq) with Seq-Well*

The Seq-Well platform was utilized as previously described[18] to capture the transcriptomes of single cells on barcoded mRNA capture beads. In brief, 10 µL of sorted $CD45^{+}$Calcein $Blue^{+}$ PBMCs were mixed 1:1 with the viability stain trypan blue and counted using a hemocytometer. The cells were resuspended in RPMI + 10% FBS at a final concentration of ~100,000 cells/mL, and 20,000-25,000 cells in 200 µL were added to each Seq-Well array preloaded with barcoded mRNA capture beads (ChemGenes). Two arrays were used for each sample to increase cell numbers. The arrays were then sealed with a polycarbonate membrane (pore size: 0.01 µm), cells were lysed, transcripts were hybridized to the beads, and the barcoded mRNA capture beads were recovered and pooled for reverse transcription using Maxima H-RT (Thermo Fisher EPO0753), and all subsequent steps. After an Exonuclease I treatment (NEB M0293L) to remove excess primers, whole transcriptome amplification (WTA) was carried out using KAPA HiFi PCR Mastermix (Kapa Biosystems KK2602) with 2,000 beads per 50 µL reaction volume. Libraries were then pooled in sets of eight (totaling 16,000 beads) and purified using Agencourt AMPure XP beads (Beckman Coulter, A63881) by a 0.6X SPRI followed by a 0.8X SPRI and quantified using Qubit hsDNA Assay (Thermo Fisher Q32854). Quality of the WTA product was assessed using the Agilent hsD5000 Screen Tape System (Agilent Genomics) with an expected peak >800 bp tailing off to beyond 3000 bp, and a small/non-existent primer peak, indicating a successful preparation. Libraries were then constructed using a Nextera XT DNA library preparation kit (Illumina FC-131-1096) on a total of 750 pg of pooled cDNA library from 16,000 recovered beads using index primers as previously described[18]. Tagmented and amplified sequences were purified using a 0.8X SPRI ratio yielding library sizes with an average distribution of 500-750 bp in length as determined using an Agilent hsD1000 Screen Tape System (Agilent Genomics). Two Seq-Well arrays were sequenced per NextSeq500 sequencing run with an Illumina 75 Cycle NextSeq500/550 v2 kit (Illumina FC-404-2005) at a final concentration of 2.4 pM. The read structure was paired end with Read 1, starting from a custom read 1 primer, covering 20 bases inclusive of a 12-bp cell barcode and 8-bp unique molecular identifier (UMI), then an 8-bp index read, and finally Read 2 containing 50 bases of transcript sequence.

*Seq-Well alignment, cell identification, cell type separation*

Read alignment, cell barcode discrimination and UMI/transcript collation were performed as in Ordovas-Montanes et al[8] using a hg19 reference. Initially, we aligned the sequences from P1 to a combined HIV + hg19 genome using the consensus sequence of HIV clade C viruses from the HIV Sequence Database (https://www.hiv.lanl.gov/content/sequence/HIV/mainpage.html). After alignment, however, we measured 0-2 cells with HIV transcript alignments per array; therefore, we used the standard hg19 reference for our analysis. UMI collapsed data was used as input into Seurat[102] (version 2.3.4) for cell and gene trimming, and downstream analysis. The following steps were performed on all of the arrays processed from a single individual, on an individual-by-individual basis. Any cell with fewer than 750 UMIs or greater than 6,000 UMIs (0-5 cells/array), and any gene expressed in fewer than 5 cells were discarded from downstream analysis. This cells-by-genes matrix was then used to create a Seurat object for each individual. Cells with > 20% of UMIs mapping to mitochondrial genes were then removed (50-100 cells/array). These objects (one per individual) were then merged into one object for pre-processing and cell-type identification

The combined Seurat object was log-normalized with a size factor of 10,000, and scaled without centering. Additionally, linear regression was performed to remove unwanted variation due to cellular complexity (nUMI) and low-quality cells (percent.mito). Subsequently, 3,251 variable genes were identified using the "LogVMR" function, and the following cutoffs: x.low.cutoff=0.01, x.high.cutoff=10, y.cutoff=0.25. Principal Component Analysis (PCA) was performed over these genes, and the top 17 PCs were chosen for clustering and embedding based on the curve of variance described by each PC and the genes most contributing to each PC. Next, FindClusters (SNN graph + modularity optimization) with resolution = 0.5 was used to generate 13 clusters, and the Fourier transform tSNE implementation[103] with 2,000 iterations to embed the data into 2-dimensional space.

Cluster identity was assigned by finding differentially expressed genes using Seurat's implemented Wilcoxon rank sum test, and then comparing those cluster-specific genes to previously published datasets[18-21]. One cluster exhibited no cluster-specific genes; the cells from this cluster were embedded centrally in the tSNE, and upon further investigation expressed both myeloid and lymphocyte markers. Therefore, these cells were removed as multiplets (when multiple cells enter the same well in the Seq-Well array). After multiplet removal, 65,842 cells were captured across all samples processed. The remaining 12 clusters included subsets of major circulating immune cells (see Table S2 for marker genes). These clusters were merged by parent cell type (T cell, cytotoxic T cell, B cell, plasmablast, DC, monocyte) for downstream analysis, as variation in the SNN graph parameters weakly affected cluster assignment to the subsets.

As NK cells share many markers transcriptionally with cytotoxic T cells[21], clustering in our data set did not separate these two cytotoxic cell types. NK cells were annotated based on expression of CD3 (*CD3D*, *CD3E*, *CD3G*), CD16 (*FCRG3A*), and KLRF1. CD56 (*NCAM*) was not highly expressed in our data, and therefore was not used to separate NK cells. Any cell with a cluster identity belonging to the cytotoxic T cell cluster that lacked CD3 expression or expressed CD16/KLRF1 was annotated as an NK cell. With this annotation, we noted distinct transcriptional responses between NK cells and CTLs both as a function of time and gene membership (Fig. 2C, Fig. 3C-E).

For downstream analysis of temporal variation in expression, the dataset was separated by individual and cell type: CD4+ T cells, NK cells, CTLs, proliferating T cells, B cells, plasmablasts, mDCs, and monocytes.

#### *Cell Type Normalization*

Once separated by cell type and individual, the single-cell transcriptomes were processed on a cell-type by cell-type basis across all time points. For each cell type, the presence of residual contaminant RNA or doublets was assayed by scoring every cell against a set of contaminant genes from other cell types built from our marker list used to discern cluster identity (see Table S8 for cell-type specific contaminant gene lists and cut-offs). Cells with high contamination scores (0-10% of cells) were subsequently removed from further analysis to avoid unwanted variation in the subsequent unsupervised module discovery. Following contamination filtering, the data underwent scaling and normalization, followed by variable gene discovery (~400-1,000 genes, dependent on cell type and cell number). PCA was then applied on these limited set of genes, followed by projection to the rest of the genes in the dataset.

#### *Module Discovery*

For the module analysis, we subset our data on the top and bottom 50 genes, after projection, for the first 3-9 PCs (dependent on the variance described by each PC, and genes contributing to each PC) as input for WGCNA functions[27]. Following the WGCNA tutorial (https://horvath.genetics.ucla.edu/html/CoexpressionNetwork/Rpackages/WGCNA/Tutorials/), an appropriate soft power threshold was chosen to calculate the adjacency matrix. As scRNA-seq data is impacted by transcript drop-out (failed capture events), adjacency matrices with high power further inflate the impact of this technical limitation, and yield few correlated modules. Therefore, when possible, a power was chosen as suggested by the authors of WGCNA (i.e., the first power with a scale free topology above 0.8); however, in instances where this power yielded few modules (fewer than 3), we decreased our power. Next, an adjacency matrix was generated using the selected soft power, and it was transformed into a Topological Overlap Matrix (TOM). Subsequently, this TOM was hierarchically clustered and the cutreeDynamic function with method "tree" was used to generate modules of correlated genes (minimum module size = 10). Similar modules were then merged using a dissimilarity threshold of 0.5 (i.e., a correlation of 0.5).

To test the significance of the correlation structure of a given module, a permutation test was implemented. Binning genes in the true module by average gene expression (# bins = 10), genes with the same distribution of average expression from the total list of genes used for module discovery were randomly picked 10,000 times. For each of these random modules, a one-sided Mann-Whitney U test was performed to compare the distribution of dissimilarity values between the genes in the true module and the distribution of dissimilarity values between the genes in the random module. Correcting the resulting p-values for multiple hypothesis testing by Benjamini-Hochberg FDR correction, a module was considered significant if fewer than 500 tests ($p < 0.05$) had FDR > 0.05.

Since we were interested in identifying modules of genes that changed in expression as a function of time, another permutation test was implemented to identify modules that significantly vary from pre-infection. First, every cell was scored for the genes within the module, using the AddModuleScore function in Seurat. As testing for differences in distribution is sensitive to sample number, a sample size (***s***) was selected based on the number of cells present at any given time point within a cell type. The smallest ***s*** used was 10; this cutoff was chosen based on the least frequent cell types having ~100 cells total across all time points within an individual. If a time-point had fewer than 10 cells, that point was not used in the testing. In the case of plasmablasts and mDCs in multiple individuals, more than three time points had fewer than 10 cells, and therefore no modules were considered significantly variant in time. To determine if module expression varied over time, 1,000 Mann-Whitney-U tests between the distribution of scores from ***s*** random cells at pre-infection and ***s*** random cells from each other time point were performed. For each time point, the p-values from the 1,000 tests were then averaged. After FDR correction, if $q < 0.05$ for any time point, the module was considered to significantly vary in expression in time. Our approach and tests have been written as functions in R and have been included in the Supplementary Files (available upon request).

*Module Grouping and Gene Set Analysis*

In order to more easily compare modules by temporal pattern within and between individuals, fuzzy c-means clustering was applied to all of the modules in a given individual using the Mfuzz package[60] (version 2.38.0). We chose to use fuzzy c-means clustering to allow us to understand the extent of membership of a given module to its assigned cluster. For each individual, c was chosen to be 5-7 such that diverse temporal patterns were separated, minimizing the number of clusters containing fewer than 3 modules. These groupings of modules were then annotated by similar scoring patterns across patients, taking into consideration that infection time is not the same for every individual (Fig. S5).

Gene Set Analysis on modules was performed using Ingenuity Pathway Analysis (IPA, Qiagen Inc.). Only gene names were supplied for analysis, and submitted for Core Analysis with the Experimentally Observed confidence setting. In Fig. 3A-E, the pathways annotated were taken from either the Canonical Pathways or Diseases & Functions results. For the upstream driver analysis in Fig. 3F-G, upstream drivers were selected by the following criteria: significant ($p < 0.001$) in at least two modules of any given cell type, with at least 5 genes in the gene set. As the gene sets annotated in IPA are quite large and share many genes, the edges in our network were restricted to only those upstream drivers who shared 3 or more genes. To achieve finer grain temporal resolution on putative inducers of immune response, the union of enriched genes for each upstream driver from modules within a given cell type was used to generate scores against the single-cell expression data. Only upstream driver scores that demonstrated temporal variability (as described above) were included. We report the median scores at each time point for each upstream driver.

The gene set enrichment analysis in Fig. 4 was performed using parts of MSigDB v6.2[104,105] (http://software.broadinstitute.org/gsea/msigdb). Multiple hypothesis testing was corrected by the Benjamini-Hochberg FDR procedure. The specific collections of gene sets used are reported in the figure legends.

*scRNA-seq of pDCs with SMART-Seq2 and analysis*

Reverse transcription, WTA, and library preparation of single pDCs in 96-well plates was performed as previously described[58]. Samples were sequenced on an Illumina NextSeq 500/550 instrument with an Illumina 75 Cycle NextSeq500/550 v2 kit (Illumina FC-404-2005) using 30-bp paired-end reads. Given difficulties acquiring pDCs from pre-infection samples due to limited cell numbers, we sequenced pDCs from the peak interferon response and the 1-year time points in each individual. Reads ($5*10^5$ – $3*10^6$/cell) were aligned to the hg38 (GENCODE v21) transcriptome and genome using RSEM[106] and Tophat[107], respectively. After trimming low quality cells (cells with <25,000 mapped reads or <1,000 genes), the remaining cells had a median of 122,000 mapped reads and 2,866 genes. Pre-processing and differential expression analysis were conducted in Seurat[102] using the Wilcoxon rank sum test. To test for differences in IFN responsiveness, individual-specific IFN response gene lists were used to generate scores in the pDCs using the AddModuleScore function in Seurat. The gene list used to score in each individual was chosen by including any gene that appeared at least twice in the modules that belonged to MM3 for that individual (see Fig. S4D).

#### *Luminex Cytokine Measurements*

Matching plasma cytokine levels were determined in duplicate using a multiplexed magnetic bead assay (Catalogue number: LHC6003M, Life Technologies) in accordance with the manufacturer's instructions. Briefly, a mixture of beads that were coated with anti-cytokine antibodies were prewashed and then incubated with the plasma samples. They were then co-incubated with a mixture of biotinylated detector antibodies followed by R-phycoerythrin (R-PE) conjugated streptavidin. A magnetic separator was used to wash the beads between incubations. Fluorescence intensity was determined on a Bio-Plex 200 system. Concentrations of the cytokines in the samples were then determined by interpolating on sigmoid 4-parameter logistic regression standard curves.

#### *Other Statistical Methods*

To determine TRBV or TRAV overabundance, we performed a $\chi^2$ test with Yates continuity correction. This test was performed independently for TRBV and TRAV genes, taking the random sampling (scaled by transcript detection) to be:

$$\frac{1}{\#\text{ of detected TRAV or TRBV genes}} * \frac{\#\text{ total alignments to TRAV or TRBV genes}}{\#\text{ of detected TRAV or TRBV genes}}$$

with # detected TRBV genes = 24; # detected TRAV genes = 35; # total alignments to TRBV genes = 379; # total alignments to TRAV genes = 525.

### SUPPLEMENTARY TEXT

*Interferon Production by pDCs*

While pDCs recognize HIV RNA by TLR7 and are thought to be the instigators of the IFN response[108], the timescale and intensity of IFN production by these cells after initial sensing has yet to be explored. Moreover, pDCs are known to home to lymph nodes in acute SIV infection[109]; therefore, the lack of detection of IFN may be in part due to sampling site. Our data suggest that a more complete temporal characterization in lymphoid tissues will be needed to appreciate their role during hyper-acute HIV infection.

*Non-overlapping B cell modules in MM1*

While B cell modules were present in two individuals (P1 & P2) in MM1, they actually expressed divergent gene expression patterns (Fig. S6B): B cells from P1 upregulated *IGHM*, *CXCR4*, and *IL4R*, genes associated with naïve B cells[110,111]; B cells from P2, meanwhile, upregulated mitochondrial genes, a potential sign of cellular stress (M4). Peaking at 6 months post-HIV detection, P2 also upregulated *IGHG1-4*, *CD52*, and *HLA-DRA* (M2), genes indicative of mature, class-switched cells; P1 demonstrated a similar module in time and gene membership (M1) of these B cells, but this module clustered into MM5 in this individual. HIV has previously been shown to induce B-cell dysfunction[112] and other work in the FRESH study has measured high variability amongst B-cell phenotypes during acute infection[113]. Given the success of NHP S(H)IV vaccine models[114,115], it may be possible to further interrogate the phenotype and function of B cells, in addition to their antibody production capacity, during hyper-acute infection in these models.

### SUPPLEMENTARY FIGURES

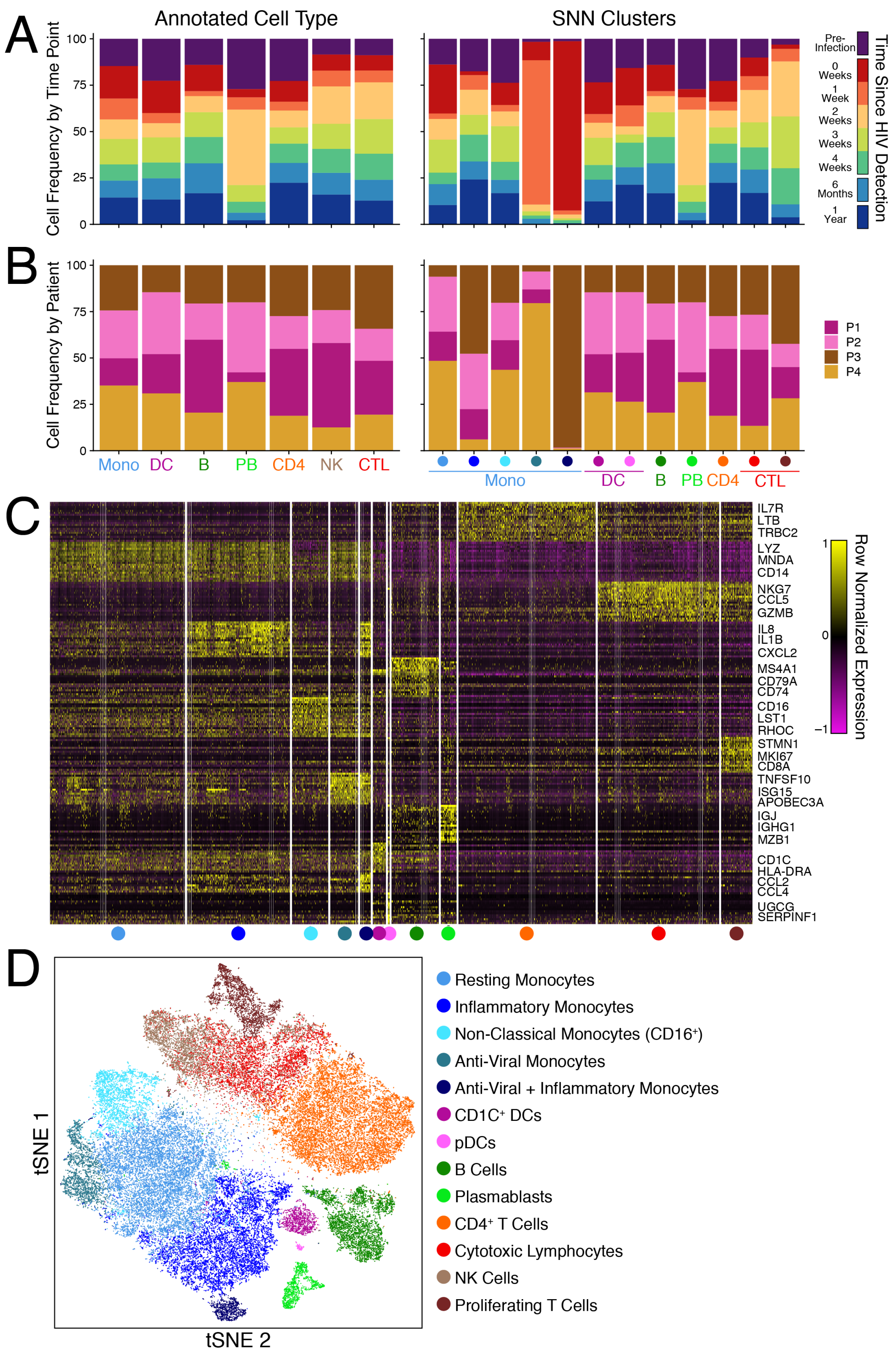


**Fig. S1: Patient and time point breakdown by cluster and cluster annotation.** (**A**) Time point and (**B**) patient cell frequency by annotated cell type (left) and shared-nearest neighbors (SNN) clusters (right). (**C**) Heatmap of the top 10 genes differentiating each SNN cluster (Wilcoxon rank sum test). (**D**) tSNE embedding of dataset colored by SNN cluster and annotated based on genes in **C** and Table S2.

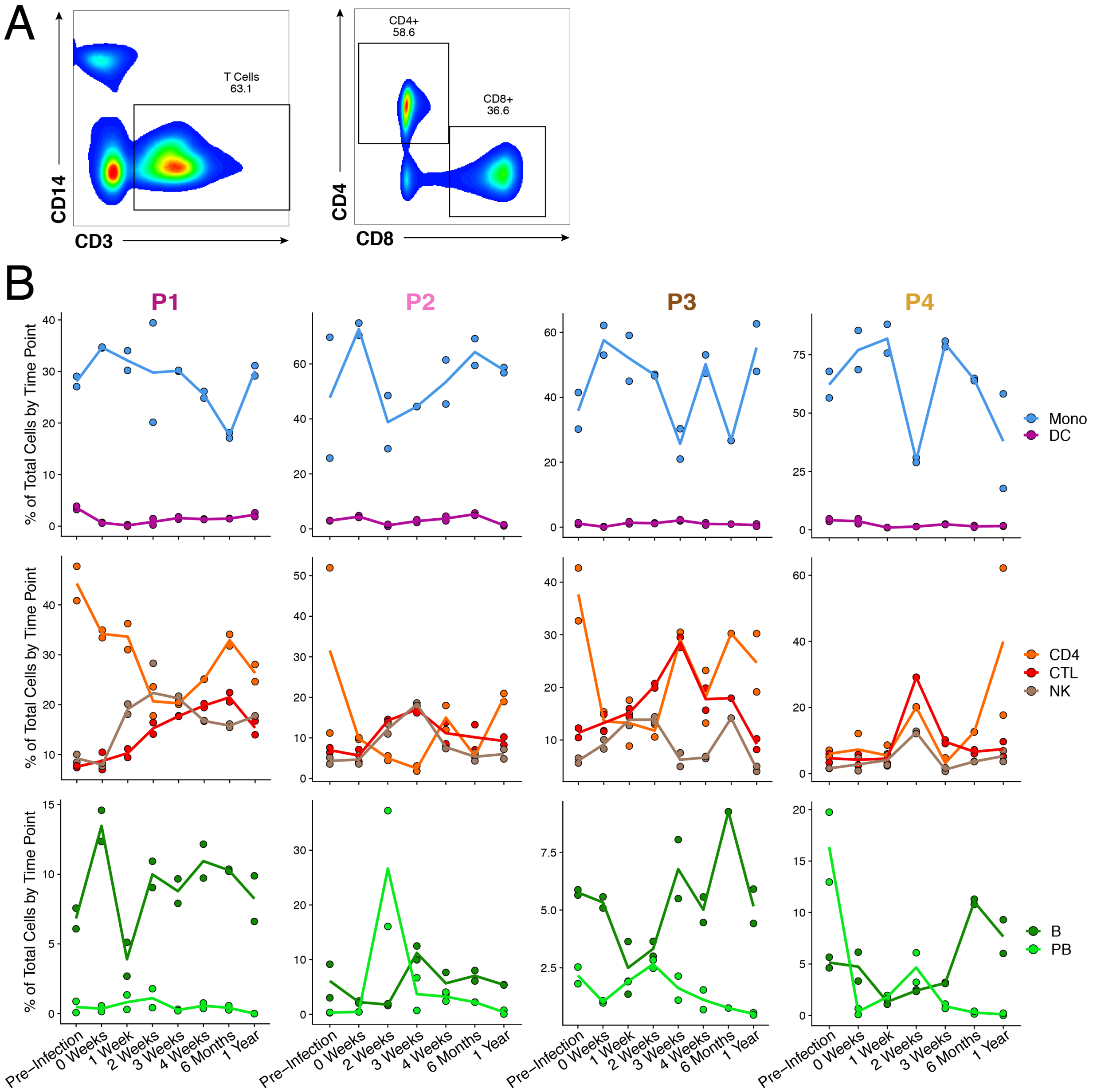


**Fig. S2: Cell frequency by individual and cell type.** (**A**) Representative gating scheme for CD4+ and CD8+ T cells. (**B**) Cell type frequency calculated from total cells measured within an array. Lines represent average between duplicate arrays. Columns are separated by individual.

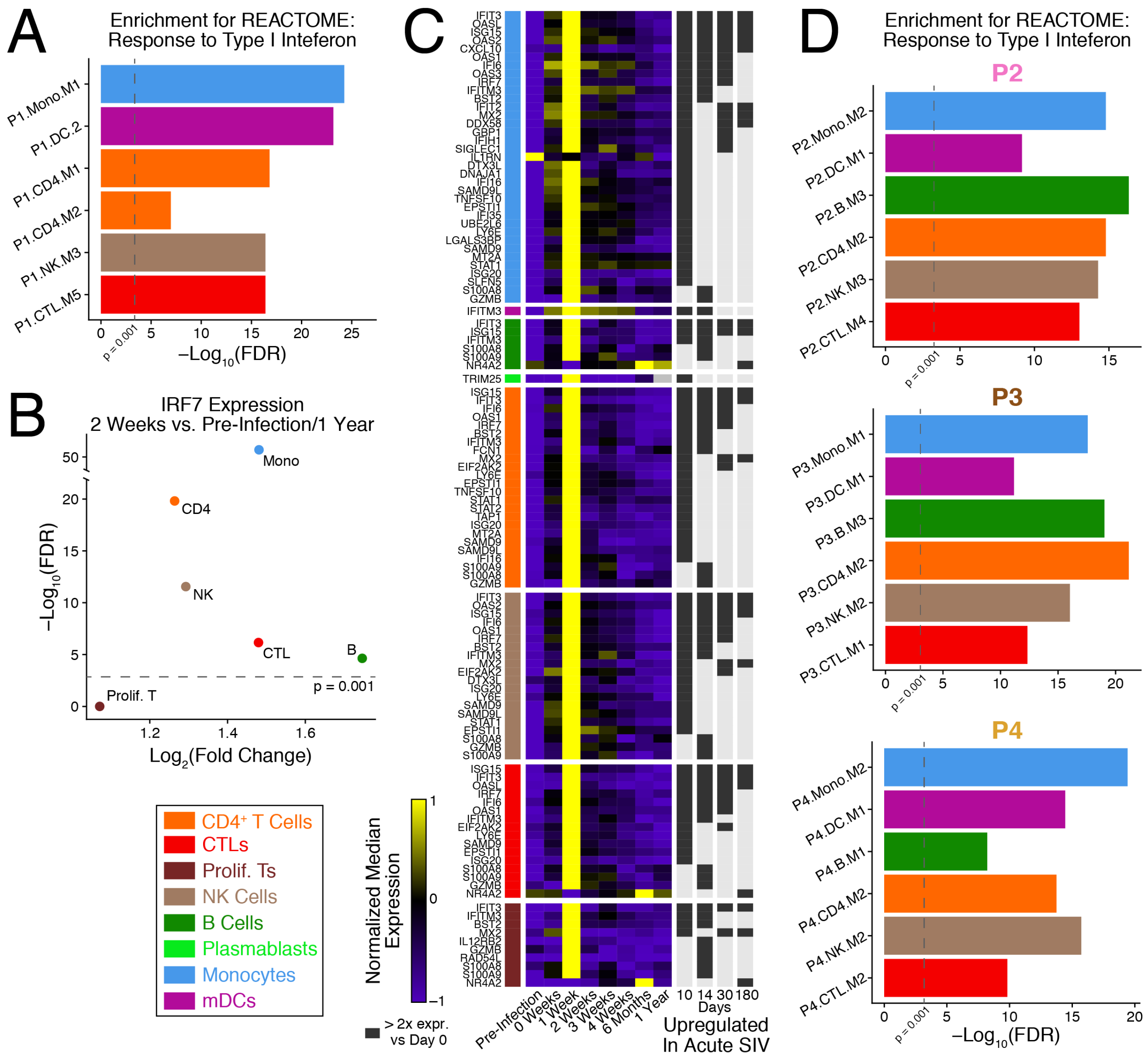


**Fig. S3: Gene modules that align near peak viremia are enriched for response to interferon and match responses observed in acute SIV infection in rhesus macaques.** (**A**) Enrichment of modules from P1 in Fig. 2B against the REACTOME: Response to Type I Interferon gene set; FDR corrected hypergeometric test. (**B**) Differential expression results for *IRF7* in each cell type (except plasmablasts and mDCs which do not have enough cells to test, n < 4) between cells from 2 weeks and pre-infection + 1 year; implemented using the "bimod" likelihood ratio test in Seurat. (**C**) Median expression of genes upregulated in SIV infection of rhesus macaques compared to day 0 (fold change > 2) in Bosinger et al. (*47*). (**D**) Same as **A** for the modules in Fig. 2D for P2, P3, and P4.

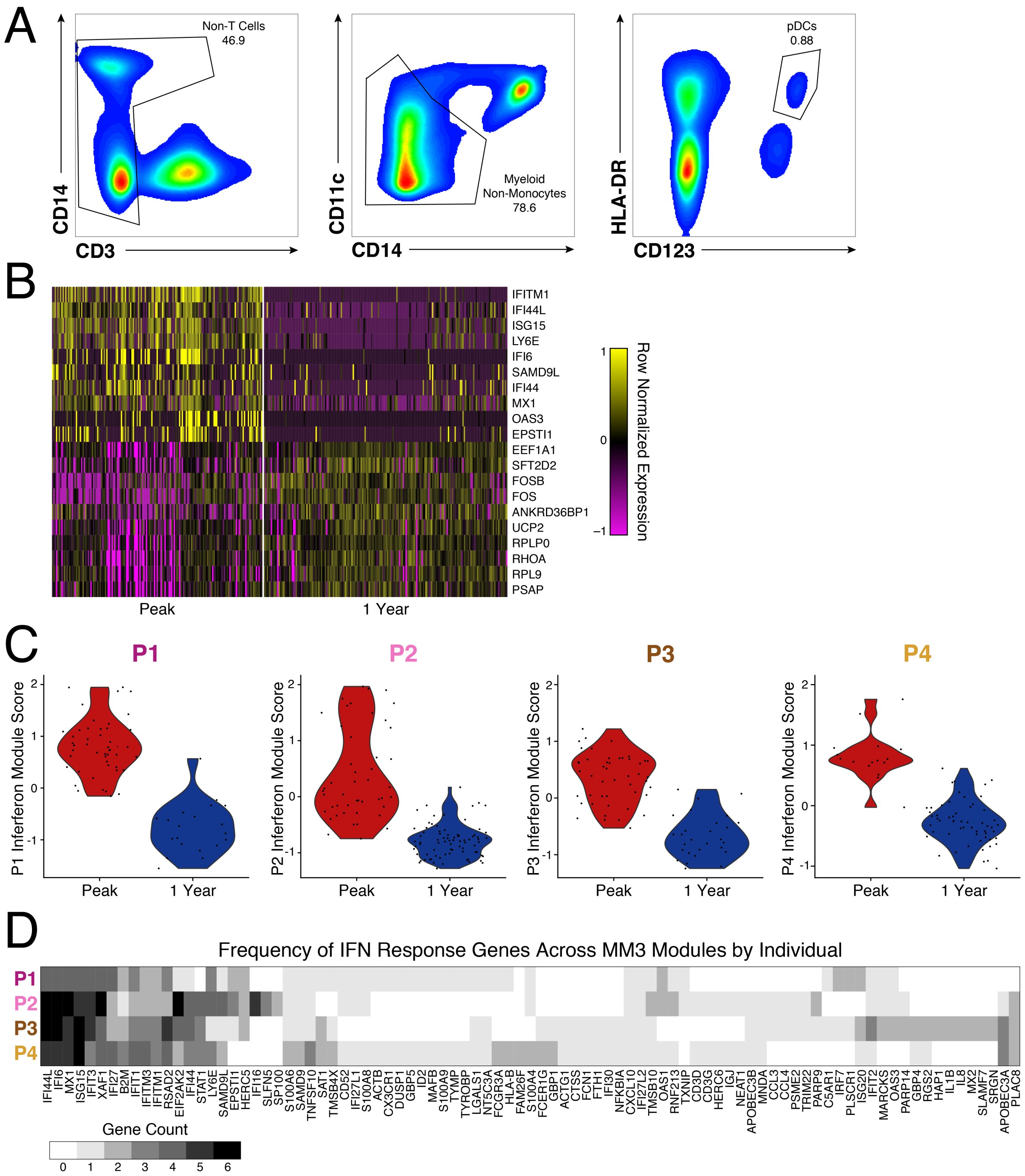


**Fig. S4: Plasmacytoid Dendritic Cells (pDCs) demonstrate similar interferon responses at the same time as other cell types.** (**A**) Representative gating scheme for single-cell pDC sorts. (**B**) Heatmap of genes differentially expressed between pDCs captured at the same time points as peak interferon responses and 1-year post HIV infection; implemented using a Wilcoxon rank sum test. (**C**) Scoring of pDCs in each individual using a core interferon signature specific to that individual. (**D**) Heatmap of gene frequency across interferon response modules in each individual.

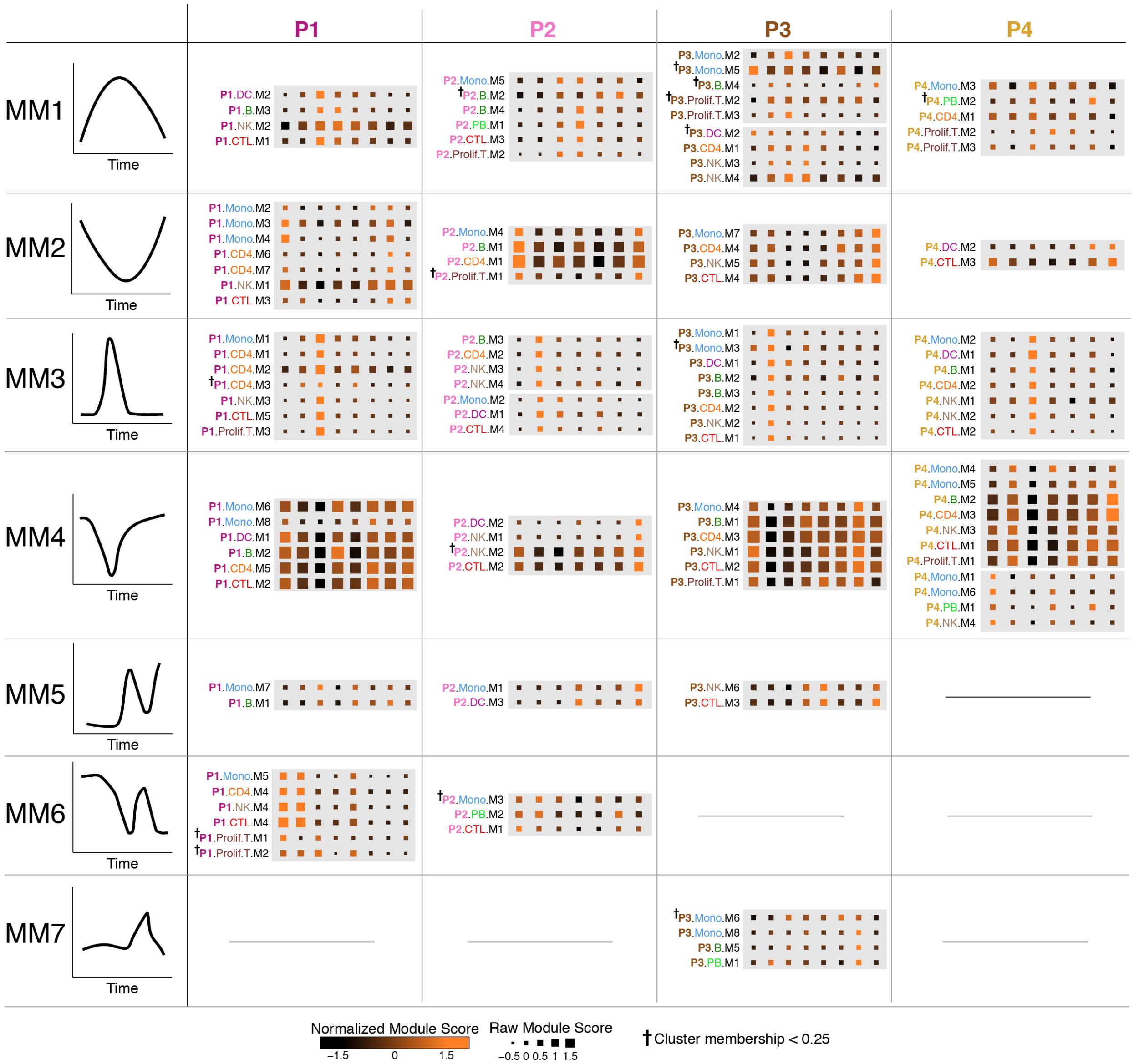


**Fig. S5: All significant temporally variant modules in all individuals grouped by fuzzy c-means clustering.** Modules grouped by fuzzy c-means clustering (see **Methods** for choice of c) reside in the same gray box. Each group of modules, or meta module (MM), were then aligned across patients based on overall temporal trend (left column). Some individuals had multiple MM with similar temporal dynamics and were grouped within the same MM. Since fuzzy c-means clustering assigns membership values to each member of a cluster, we report any modules that demonstrated low cluster membership with †.

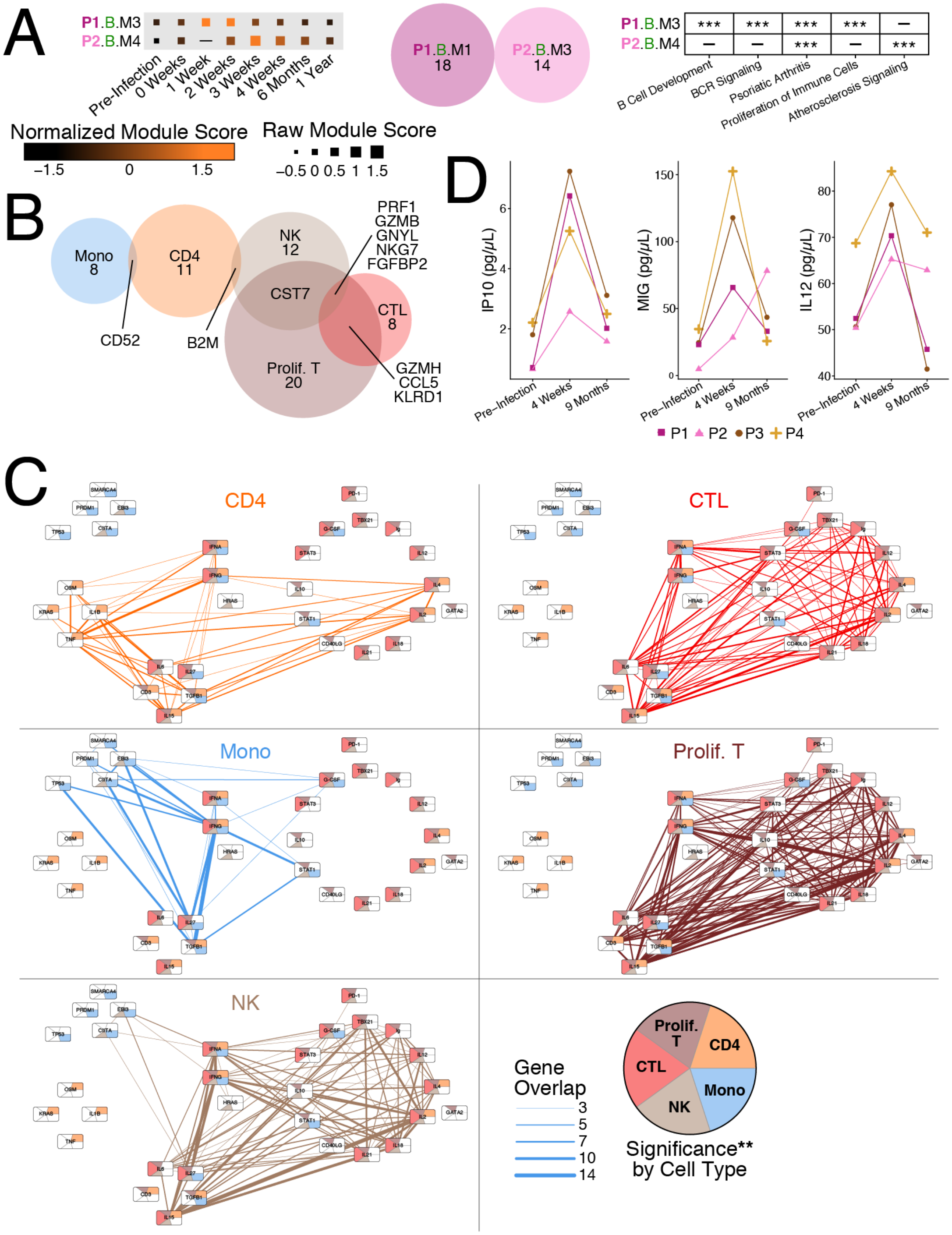

323

324 **Fig. S6: Sustained B cell modules and shared genes and upstream drivers between**
325 **individuals.** (**A**) B cell modules in MM1 with high cluster membership. (**B**) Euler diagram of
326 conserved overlapping genes between cell types from Fig. 3A-E, see Table S5. (**C**) Fig. 3F
327 displayed with only the edges from a given cell type. (**D**) Luminex measurements of IP10 (left),

MIG (center), and IL-12 (right) in matching plasma samples. Points are averages of duplicate measurements.

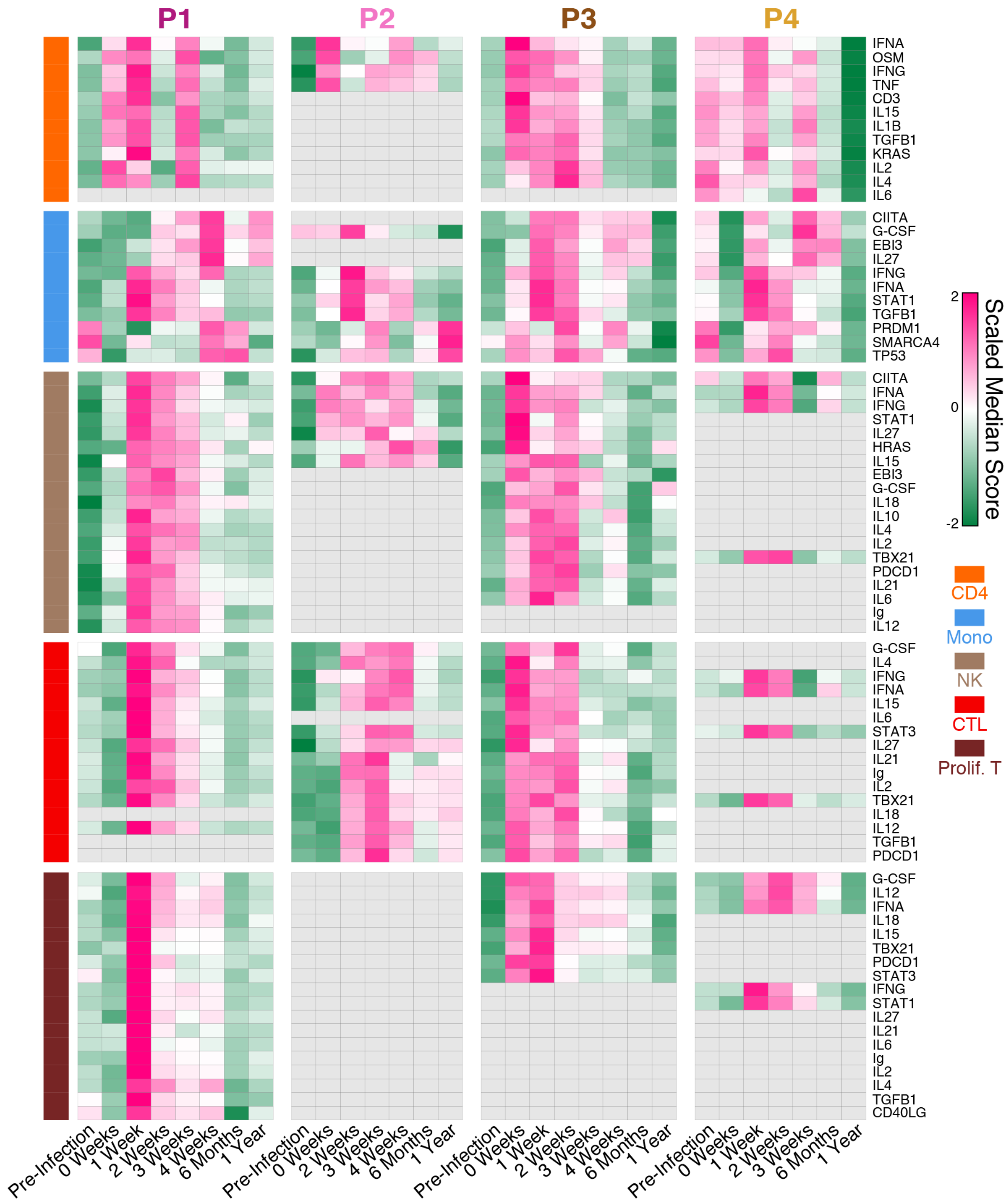


**Fig. S7: Putative upstream driver scores highlight variable response dynamics across cell types.** Median gene set scores for significantly temporally variant ($p < 0.05$) upstream drivers in all individuals. Gray boxes indicate that the upstream driver was not significantly variant in that cell type and individual.

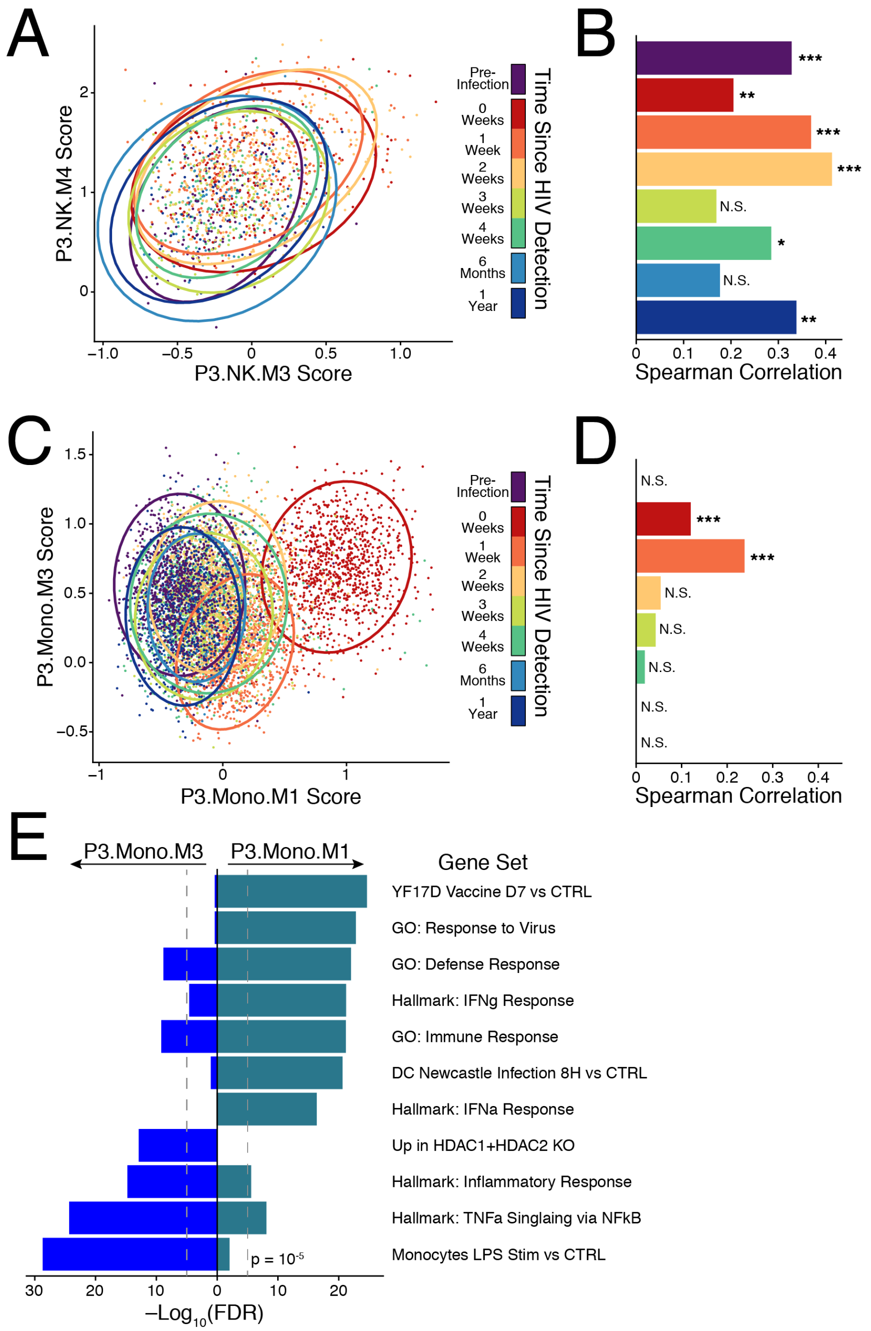


**Fig. S8: Two cases of similar temporal modules: variable correlation and variable co-expression.** (**A**) Module scores in NK cells for NK M3 and NK M4 in P3. Ellipses drawn at 95% confidence interval for cells from each time point. (**B**) Correlation (spearman's rho) between the scores for NK M3 and NK M4 at each time point. FDR corrected q-value: N.S = not significant; *q < 0.05; ** q < 0.01, *** q < 0.001. (**C & D**) Same as in **A** & **B** but for monocyte modules Mono M1 and Mono M3 in P3. (**E**) Gene set enrichment analysis of the genes in Mono M1 and Mono M3 against the following MSigDB collections: Hallmark, C2, C3, C5, and C7. FDR corrected hypergeometric test.

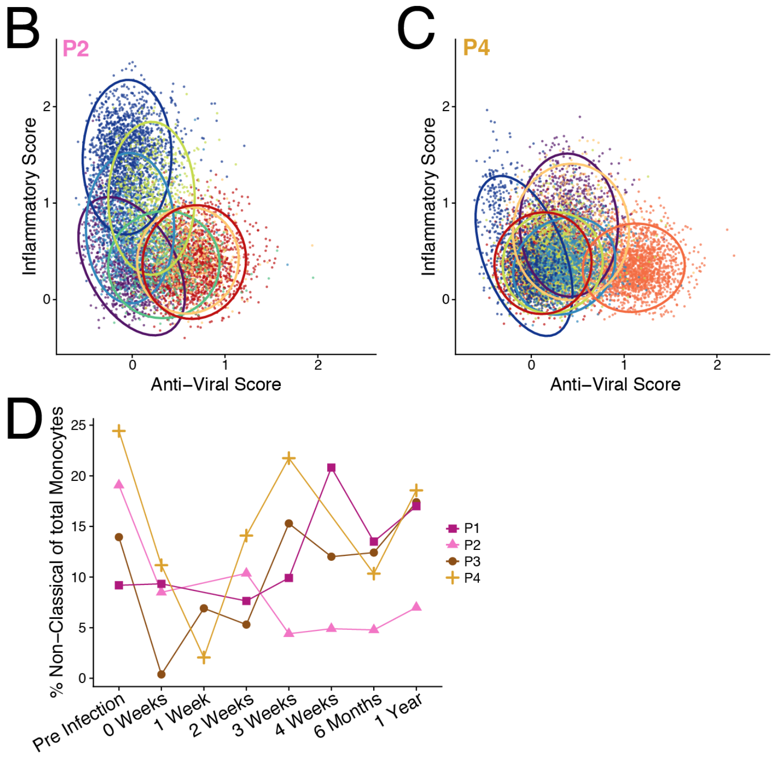


**Fig. S9: Core anti-viral, inflammatory, and non-classical programs in monocytes.** (**A**) The genes shared between individuals (present in at least two modules) that make up the inflammatory and anti-viral scores used in Fig. 4A, as well as in **B** and **C** of this figure (available upon request). (**B** & **C**) Inflammatory and anti-viral scores of monocytes by time point in P2 (**B**) and P4 (**C**). Ellipses drawn at 95% confidence interval for cells from each time point. (**D**) Percent of Non-Classical ($CD16^{+}$) monocytes of total monocytes as a function of time in each individual. Percentage calculated from cluster assignment (see Fig. S1D).

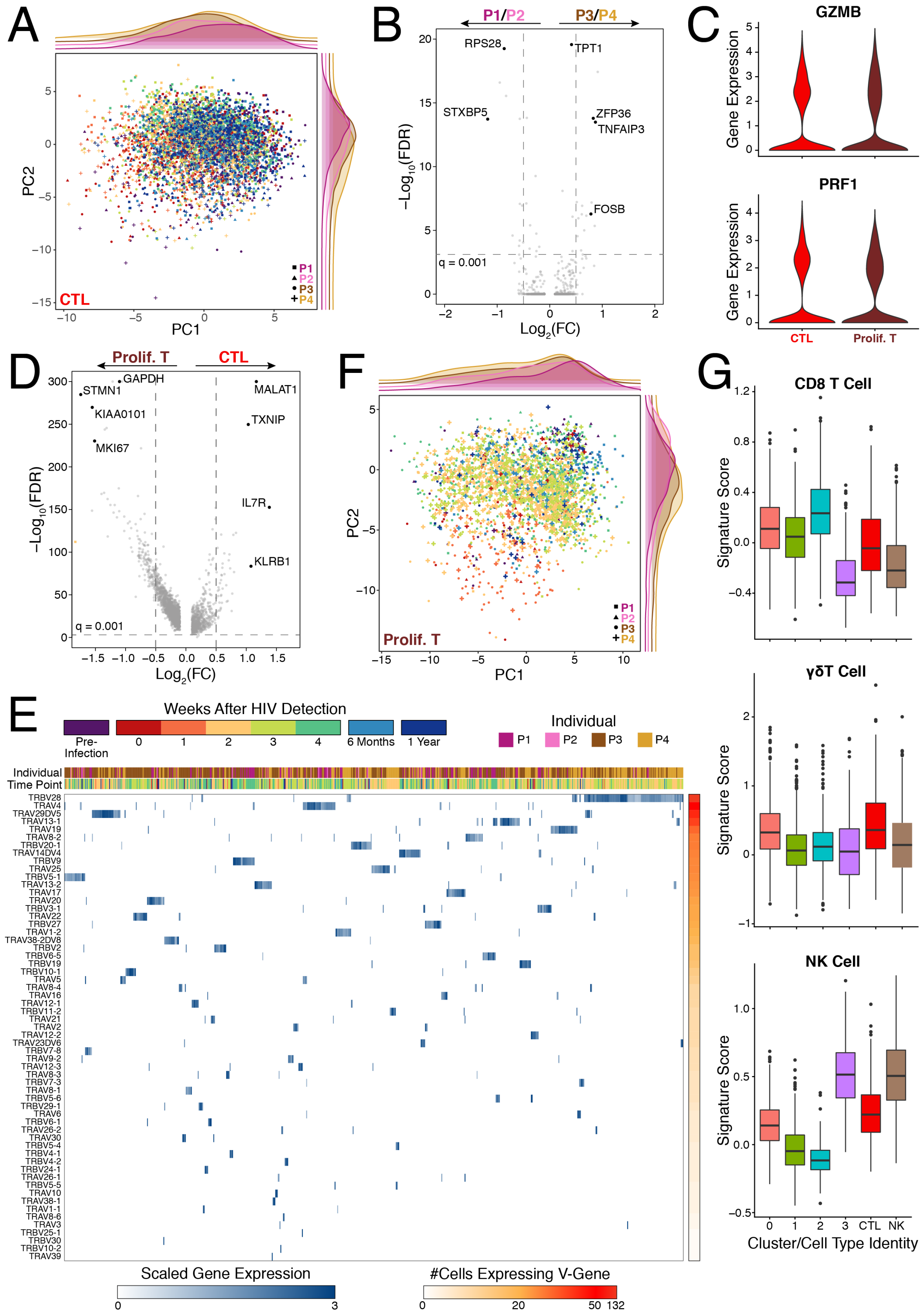
A
CTL
PC1
PC2
P1
P2
P3
P4
B
P1/P2
P3/P4
RPS28
TPT1
STXBP5
ZFP36
TNFAIP3
FOSB
q = 0.001
−Log10(FDR)
Log2(FC)
C
GZMB
PRF1
Gene Expression
CTL
Prolif. T
D
Prolif. T
CTL
GAPDH
STMN1
MALAT1
KIAA0101
TXNIP
MKI67
IL7R
KLRB1
q = 0.001
−Log10(FDR)
Log2(FC)
F
Prolif. T
PC1
PC2
P1
P2
P3
P4
G
CD8 T Cell
γδT Cell
NK Cell
Signature Score
Cluster/Cell Type Identity
0
1
2
3
CTL
NK
E
Weeks After HIV Detection
Pre-Infection
6 Months
1 Year
Individual
P1
P2
P3
P4
Individual
Time Point
Scaled Gene Expression
#Cells Expressing V-Gene

**Fig. S10: Non-proliferating and proliferating cytotoxic T cells.** (**A**) Principal component analysis of non-proliferating CTLs with patient density annotated along PC1 and PC2. (**B**) Volcano plot of differentially expressed genes between the individuals who control (P3/P4) and those who do not (P1/P2); implemented using a Wilcoxon rank sum test. (**C**) Expression of *GZMB* and *PRF1* in all CTLs and proliferating T cells. (**D**) Volcano plot of differentially expressed genes between non-proliferating CTLs and proliferating T cells; implemented using a Wilcoxon rank sum test. (**E**) Heatmap of detected TCR-$\alpha$ and TCR-$\beta$ variable chain genes in proliferating T cell clusters 0 & 1. (**F**) Same as in **A** but over proliferating T cells. (**G**) CD8 T cell (top), $\gamma\delta$T cell (middle), and NK cell (bottom) scores for each proliferating T cell cluster (see **Fig 5C**), 500 randomly sampled CTLs, and 500 randomly sampled NK cells. Signatures were established from differential expression over the single-cell dataset published by Gutierrez-Arcelus et al. (*21*). See Table S7 for all differentially expressed genes and signature score gene lists.

### SUPPLEMENTARY TABLES

| Individual | Time Point (days relative to first positive viral RNA test) | | | | | | | |
|---|---|---|---|---|---|---|---|---|
| | *Pre-Infection* | *0 Weeks* | *1 Week* | *2 Weeks* | *3 Weeks* | *4 Weeks* | *6 Months* | *1 Year* |
| P1 | -70 | 3 | 10 | 17 | 24 | 31 | 168 | 330 |
| P2 | -42 | 3 | N.C. | 17 | 24 | 31 | 161 | 329 |
| P3 | -35 | 1 | 7 | 14 | 21 | 28 | 162 | 329 |
| P4 | -95 | 1 | 7 | 14 | 24 | N.C. | 164 | 413 |

| Individual | Enrollment Age | STI at HIV Detection | Feibig Stage at Detection | HLA–A | HLA–B | HLA–C |
|---|---|---|---|---|---|---|
| P1 | 24 | Yes | I | 24:02 / 29:02 | 07:02 / 44:03 | 07:01 / 07:02 |
| P2 | 21 | N/A | I | 68:01 / 68:02 | 57:02 / 58:02 | 06:02 / 18 |
| P3 | 24 | Yes | I | 02:05 / 66:01 | 14:01 / 39:10 | 8:04 / 12:03 |
| P4 | 21 | Yes | I | 01:01 / 66:01 | 39:10 / 81:01 | 12:03 / 18 |

**Table S1: Time point, clinical information, and HLA genotype for individuals studied.**

*Tables S2-S8 are available upon request.*

**Table S2: Differentially expressed genes between SNN clusters.**

**Table S3: Gene membership of discovered modules by individual.**

**Table S4: Median module scores by timepoint in each individual.**

**Table S5: Overlapping genes between modules with sustained expression in CD4+ T cells, Monocytes, CTLs, NK cells, and proliferating T cells.**

**Table S6: Ingenuity Pathway Analysis enrichment results for shared modules with sustained expression.**

**Table S7: Differentially expressed genes in CTLs and proliferating CTLs.**

**Table S8: Genes used to determine residual RNA contamination in each cell type.**
